# Antibody:CD47 ratio regulates macrophage phagocytosis through competitive receptor phosphorylation

**DOI:** 10.1101/2020.07.31.231779

**Authors:** Emily C. Suter, Eva M. Schmid, Erik Voets, Brian Francica, Daniel A. Fletcher

## Abstract

Cancer immunotherapies often modulate macrophage effector function by introducing either targeting antibodies that activate Fc gamma receptors or blocking antibodies that disrupt inhibitory SIRPα-CD47 engagement. Yet how these competing signals are integrated is poorly understood mechanistically, raising questions about how to effectively titrate immune responses. Here we find that macrophage phagocytic decisions are regulated by the ratio of activating ligand to inhibitory ligand on targets over a broad range of absolute molecular densities. Using endogenous as well as chimeric receptors, we show that activating:inhibitory ligand ratios of at least 10:1 are required to promote phagocytosis of model antibody-opsonized CD47-inhibited targets and that lowering this ratio reduces FcγR phosphorylation due to inhibitory phosphatases recruited to CD47-bound SIRPα. We demonstrate that ratiometric signaling is critical for phagocytosis of tumor cells and can be modified by blocking SIRPα *in vitro*, indicating that balancing targeting and blocking antibodies may be important for controlling macrophage phagocytosis in cancer immunotherapy.

## INTRODUCTION

Macrophages play a critical role in cancer immunotherapy by recognizing and destroying antibody-opsonized cells. The clinical success of therapeutic antibodies, including anti-CD20 clearance of chronic lymphocytic leukemia cells (Chu et al., 2018; VanDerMeid et al., 2018) and anti-CD38 clearance of multiple myeloma (van de Donk & Usmani, 2018), has been attributed in part to macrophage destruction of tumor cells via antibody-dependent cellular phagocytosis (ADCP). However, potent inhibitory molecules found on tumor cells, such as the ubiquitous “marker of self” CD47, can dampen the pro-phagocytic effect of therapeutic antibodies (Okazawa et al., 2005; Oldenborg et al., 2000). Interaction of macrophage receptor SIRPα with CD47 on target cells drives phosphorylation of SIRPα’s multiple immunoreceptor tyrosine-based inhibitory motifs (ITIMs), which then leads to inhibition of phagocytosis (Barclay & van den Berg, 2014). Though the SIRPα-CD47 inhibitory checkpoint is critically important for avoiding destruction of healthy cells, it is commonly hijacked by tumor cells, which can upregulate CD47 expression from 2 to 6 times normal surface expression to avoid recognition and destruction (Willingham et al., 2012).

Disrupting the SIRPα-CD47 inhibitory checkpoint has become an important immunotherapeutic target in recent years (Feng et al., 2019). Blockade strategies like those used to modulate T-cell activity are now being explored in macrophages, with antibodies that block SIRPα and CD47 intended to remove the brakes on tumor-targeted macrophage effector function. Promising clinical and mouse studies have recently shown that CD47 blockade therapies can be effective in bolstering the efficacy of therapeutic antibodies (Chao et al., 2010; Majeti et al., 2009; Theocharides et al., 2012; Weiskopf et al., 2016), though concerns remain about the potential for off-target effects. Since high concentrations of a blocking antibody are best for disrupting the SIRPα-CD47 inhibitory checkpoint but low antibody concentrations are best for minimizing off-target effects, doses must be found that are neither ineffective nor dangerous. While this can and is being done empirically in clinical trials (Feng et al., 2019), understanding the molecular mechanisms underlying macrophage decision-making is an important step towards designing effective and safe combination immunotherapies.

How do macrophages integrate conflicting activating and inhibitory signals? Recent work on macrophage FcγR activation has helped to clarify the molecular mechanisms responsible for antibody-dependent phagocytosis (Bakalar et al., 2018; Freeman et al., 2016), but how CD47 binding activates SIRPα and how SIRPα ITIMs counteract FcγR signaling remain active areas of research. Recently, Morrissey et al. demonstrated that SIRPα-CD47 binding inhibits macrophage integrin activation and that artificially stimulating integrins recovers some of the impaired phagocytosis (Morrissey & Vale, 2019). Previously, Tsai et al. showed that dephosphorylation of non-muscle myosin IIA was a key component of CD47-mediated shutdown of macrophages (Tsai & Discher, 2008). However, it remains possible that a more direct interaction between activating and inhibitory receptors could dictate decisions regarding macrophage effector function.

Here, we quantitatively evaluate macrophage phagocytic decision-making by systematically varying pro-phagocytic antibody and anti-phagocytic CD47 in a reconstituted target system. We find that the ratio of activating and inhibitory ligands, rather than their absolute number, determines macrophage phagocytosis, with an antibody:CD47 ratio of 10:1 necessary to overcome inhibition in the model system. We show that this ratio-dependent behavior is exhibited by both endogenous and chimeric receptors and that shifting this ratio drives SIRPα-mediated reduction in FcγR phosphorylation. Finally, we demonstrate that the activation:inhibition ratio is critical for tumor cell phagocytosis, indicating that control of macrophage effector function with inhibitory checkpoint immunotherapies will depend on the relative number of antigens and CD47 on target tumor cells.

## RESULTS

### Ratio of Antibody:CD47 drives phagocytosis in a reconstituted target assay

To study competition between inhibitory SIRPα and activating FcγR signals, we used reconstituted cell-like target particles consisting of silica beads coated in fluorescent supported lipid bilayers (SLBs). Briefly, Ni-NTA-conjugated lipids in the SLB enabled controlled attachment of His-tagged CD47 (inhibitory ligand), and the target particles were opsonized by binding anti-biotin IgG to biotinylated lipids (activating ligand) (Figure 1a). Importantly, the fluidity of the SLB facilitates the binding and subsequent enrichment of membrane-bound proteins on the target (Joffe, A. et al., 2020). After protein attachment, target particles were added to RAW 264.7 macrophage-like cell line and phagocytosis was measured by quantifying internalized target particle fluorescence using confocal microscopy. We found that target particles coated only in antibody were engulfed, but addition of CD47 reduced phagocytosis to background (no protein) levels, showing that this inhibitory axis can be reconstituted in our model system (Figure 1b).

**Figure 1.**
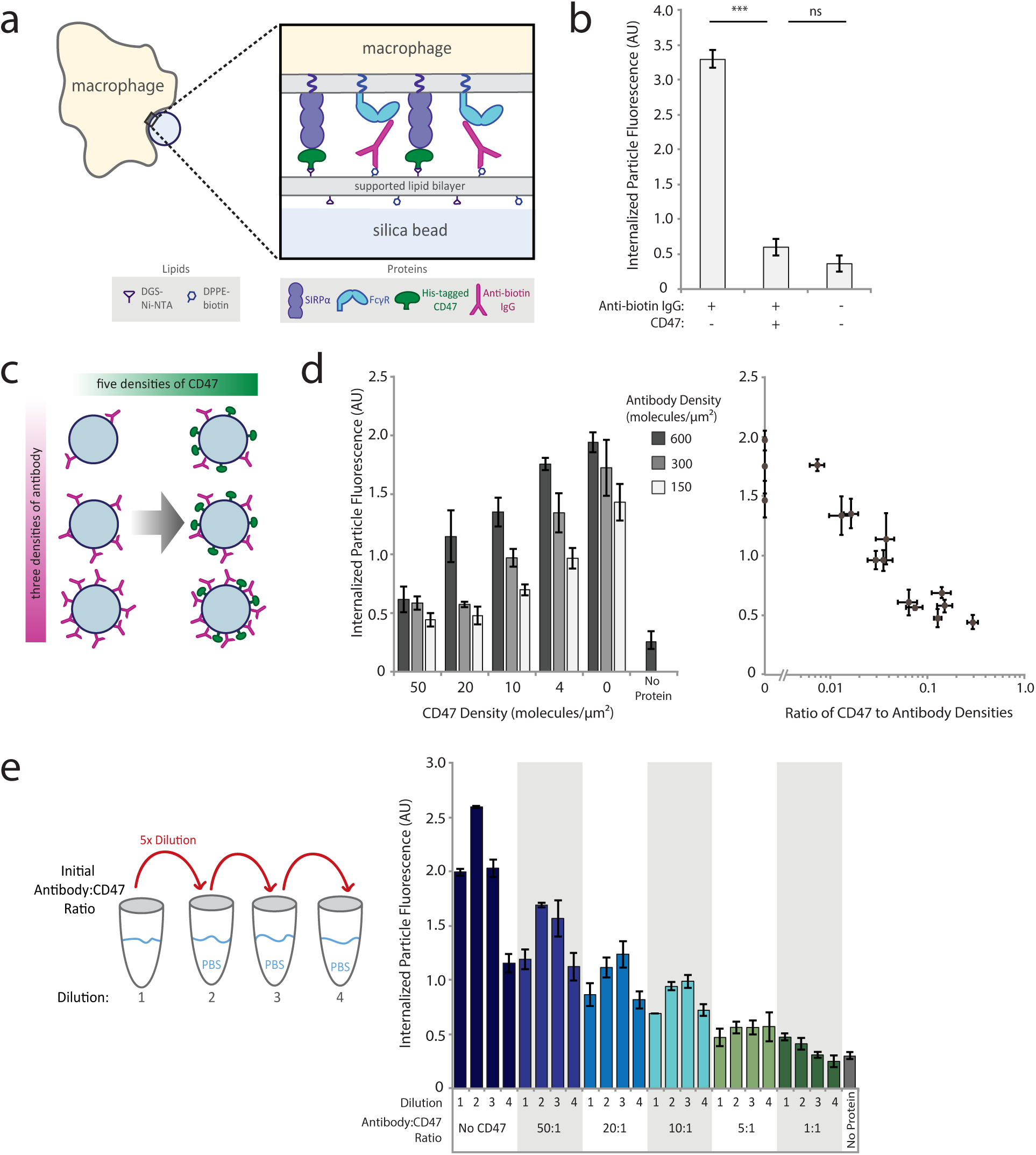
Phagocytosis is dependent on ratio of activating antibody to inhibitory CD47. (a) Schematic of experimental setup in which cell-like target particles were incubated with RAW 264.7 macrophages to assess phagocytic efficiency. (b) SLB-coated particles were incubated with 1 nM anti-biotin IgG with and without 50 nM His-tagged CD47. Quantification of average internalized target particle fluorescence shows phagocytosis of protein-coated targets as compared to empty (lipid-only) targets (right panel). Each condition is the average of three independent experiments representing a total of >300 cells. Bars represent mean ± s.e.m. (c) Systematic exploration of different anti-biotin IgG and CD47 densities on target particle surfaces. SLB-coated particles were incubated with 15 different combinations of anti-biotin IgG and CD47 concentrations, in addition to a no protein (lipid only) control. For each of three anti-biotin IgG densities (150, 300, 600 molecules/μm^2^), five CD47 densities of CD47 were added (0-50 molecules/μm^2^). (d) Internalized target particle fluorescence quantification for conditions outlined in (c) (left panel). Data replotted as a function of CD47:antibody ratio (right panel). Each condition is the average of three independent experiments representing a total of >300 cells. Bars represent mean ± s.e.m. (e) Different anti-biotin IgG to CD47 ratios were created at high concentration, then serially diluted 5-fold three times, creating a total of four concentrations for each ratio. Target particles were coated in protein dilutions, incubated with cells, and internalized particle fluorescence was quantified. Each condition is the average of three independent experiments representing a total of >300 cells. Bars represent mean ± s.e.m.

We next investigated how macrophage phagocytosis is dependent on ligand density and whether phagocytosis stops when CD47 reaches a specific threshold density on the target surface. Lipid composition of the target particle allows for orthogonal attachment of antibody and CD47, enabling us to precisely control the numbers of activating and inhibitory ligands presented to the macrophage. To verify these numbers, surface densities of fluorescently-labeled CD47 and antibody on target particles were measured using flow cytometry for every phagocytosis assay (Figure S1a). Importantly, estimated macrophage SIRPα and FcγR receptors are in excess of their corresponding ligands measured on the target particle surface.

We first measured sensitivity to ligand density by creating a panel of target particles combining three different antibody surface densities (150, 300, and 600 molecules/μm^2^), each with five different CD47 surface densities (0, 4, 10, 20, 50 molecules/μm^2^, Figure 1c). For comparison, the CD47 surface density is approximately 250 molecules/μm^2^ on red blood cells (Mouro-Chanteloup et al., 2003). Over the range of surface densities investigated, phagocytosis scaled inversely with CD47 and did not exhibit a step function at a specific CD47 density, as would be expected for threshold-governed behavior (Figure 1d). Target particles with the highest antibody density overcame CD47 inhibition better than those with lower antibody densities, indicating that phagocytosis could be responding to both CD47 and antibody densities. Indeed, by replotting the data as a function of CD47:antibody ratio, we see that phagocytosis is highly sensitive to small changes in relative CD47:antibody densities rather than absolute molecule numbers (Figure 1d).

To more directly test this ratio hypothesis, we mixed antibody and CD47 in ratios varying from 1:1 to 50:1, and then serially diluted them prior to adding proteins to SLB-coated beads (Figure 1e). This enabled us to create target particles with a fixed ratio of molecules but vary the absolute numbers of those molecules by over 100-fold (Figure S1b). We find that target particles with the same ratio of molecules yield similar levels of phagocytosis across a wide range of surface densities. We note, however, that phagocytosis can be sensitive to absolute molecule numbers when measured densities of antibody or CD47 become extremely low (<20 molecules/μm^2^ anti-biotin antibody and <2 molecules/μm^2^ CD47). For dilution series in which the first dilution shows lower phagocytosis than subsequent dilutions, we hypothesize that an excess of particle-bound and soluble antibody saturates FcγRs, effectively leading to a lower bound antibody:CD47 ligand ratio and therefore reduced phagocytosis.

### SIRPα-CD47 binding drives exclusion of CD45 and co-localizes with bound FcγRs

We next investigated how the ratio of antibody:CD47 might govern phagocytosis by studying the composition of the interface between a macrophage and target particle. To improve our ability to quantify interfacial organization and density, we used TIRF microscopy to image macrophages undergoing frustrated phagocytosis on SLB-coated glass coverslips containing the same tunable concentrations of antibody and CD47 as the target particles used above (Figure 2a).

**Figure 2.**
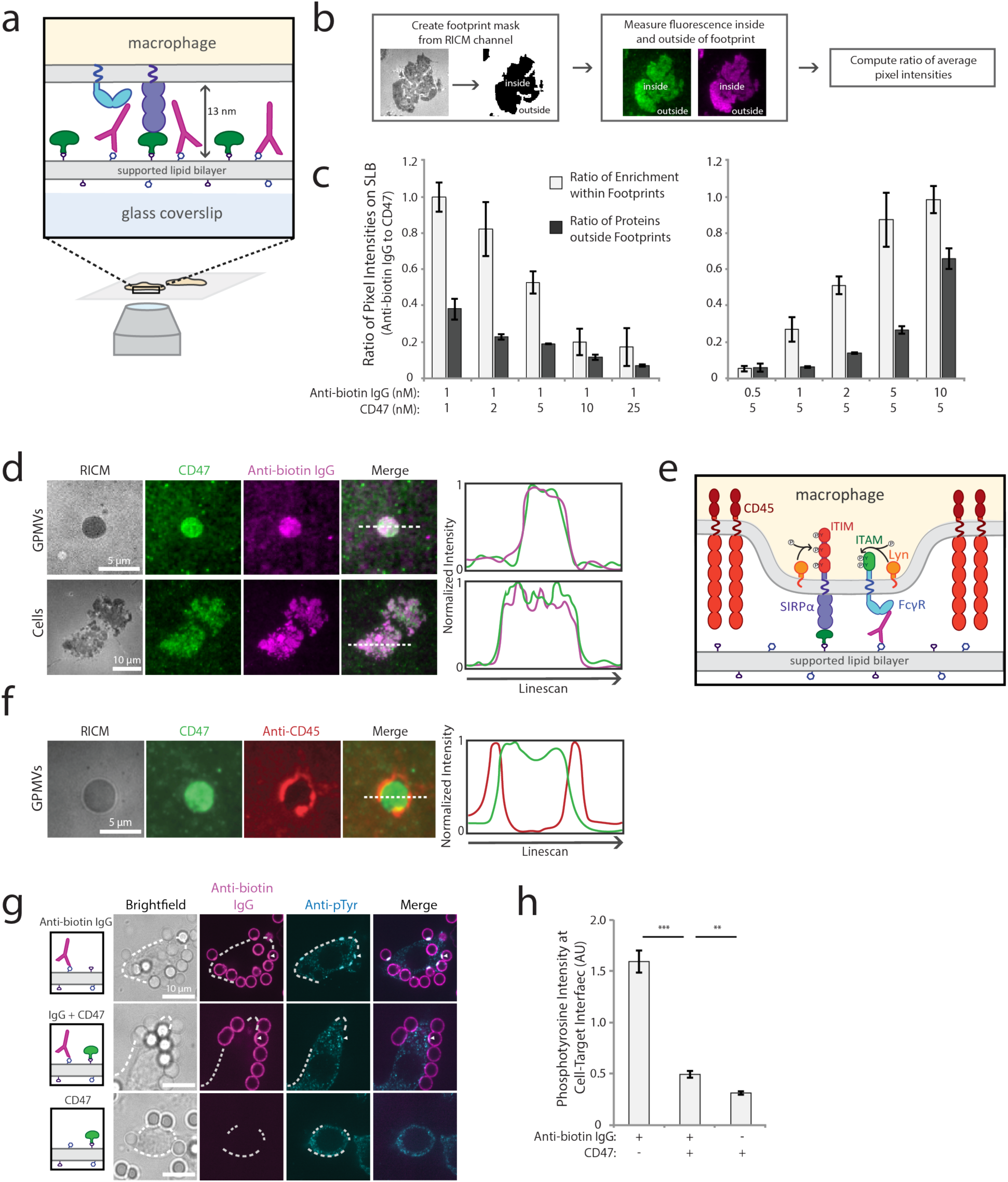
SIRPα-CD47 co-localizes with FcγR-antibody complexes and excludes CD45. (a) Schematic of TIRF experimental set-up to image RAW 264.7 macrophages on SLB. (b) Workflow to quantify enrichment of CD47 and anti-biotin IgG on an SLB within a RAW 264.7 macrophage cell footprint. SLBs with variable concentrations of AlexaFluor647-labeled anti-biotin IgG (magenta) and AlexaFluor488-labeled CD47 (green) were created, and cell footprints were identified from RICM images. Average ratio of pixel intensities (anti-biotin IgG:CD47) was quantified inside and outside footprints. (c) Average ratio of anti-biotin IgG and CD47 pixel intensities for SLBs of varying composition. Each condition is the average of three independent experiments representing a total of >200 footprints. Bars represent mean ± s.e.m. (d) Representative TIRF fluorescent images of RAW 264.7 macrophages or GPMVs generated from RAW 264.7 cells on a planar SLB. SLBs were coated with 1 nM AlexaFluor647-labeled anti-biotin IgG (magenta) and 5 nM AlexaFluor488-labeled His-tagged CD47 (green). Linescans of representative images show co-localization of anti-biotin IgG and CD47 in cell and GPMV footprints. (e) Schematic of FcγR ITAM and SIRPα ITIM phosphorylation by Lyn kinase upon CD45 phosphatase exclusion. (f) Representative TIRF fluorescent image of GPMVs from RAW 264.7 macrophages on a SLB with AlexaFluor488-labeled His-tagged CD47 (green). CD45 was labeled prior to GPMV addition to the SLB with AlexaFluor555-labeled anti-CD45 antibody (red). Linescan of representative image shows CD45 exclusion from interface. (g) Fluorescent confocal images of fixed RAW 264.7 macrophages interacting with target particles. Top row shows representative images of macrophages interacting with particles coated only in 1 nM AlexaFluor647-labeled anti-biotin IgG (magenta). Middle row shows particles coated in 1 nM AlexaFluor647-labeled anti-biotin antibody plus 50 nM CD47, and the bottom row shows particles coated only in 50 nM CD47. Cell outlines are shown by white dotted lines. Anti-phosphotyrosine was visualized through AlexaFluor546-labeled secondary antibody (cyan). (h) Quantification of anti-phosphotyrosine signal at bead-macrophage interfaces. Each condition is the average of three independent experiments representing a total of >100 cell-bead interfaces. Bars represent mean ± s.e.m.

We first investigated whether receptor engagement at the interface was proportional to the ratio of antibody:CD47 presented on the beads. We tested this by dropping cells on to planar SLBs with varying antibody:CD47 ratios and measuring enrichment within the macrophage footprint (Figure 2b). We find that average enrichment within the footprint scales with the ratio of ligands present on the SLB, indicating that binding of the two ligand-receptor pairs is independent of each other (i.e. local enrichment of one does not inhibit enrichment of the other) (Figure 2c).

Next, we examined how the size of bound SIRPα-CD47 compared with the size of antibody bound to FcγR. We previously found that macrophage phagocytosis is dependent on size-based segregation of the inhibitory phosphatase CD45 from the antibody-bound FcγR (Bakalar et al., 2018). Specifically, FcγR binding to an antibody targeting the membrane, which creates a membrane-membrane gap of approximately 12 nm (estimated from crystal structures (Bakalar et al., 2018; Lu et al., 2011)), results in efficient phagocytosis due to maximum exclusion of CD45 (Bakalar et al., 2018). This activating interface gap almost perfectly matches with the size of SIRPα-CD47 interaction (∼13 nm estimated by crystal structure (Hatherley et al., 2009)). This would allow bound SIRPα-CD47 to be remain in the gap formed by a short phagocytic interface, potentially inhibiting phagocytosis. To test this prediction, we used the SLB system and TIRF imaging to assess co-localization of the two binding pairs, FcγR-antibody and SIRPα-CD47, at the same interface. We used RAW 264.7 cells to examine receptor-ligand co-localization in live macrophages and giant plasma membrane vesicles (GPMVs) generated from RAW cells to examine receptor-ligand co-localization in a system that contains a similar plasma membrane protein composition but lacks an active cytoskeleton (Sezgin et al., 2012). In both the RAW cells and GPMVs, SIRPα-CD47 and FcγR-antibody were enriched and co-localized in the footprint, indicating that these two opposing pathways can co-exist at an interface due to their similarity in size (Figure 2d).

Size-dependent segregation of CD45 from the small gap created by FcγR engagement with antibodies is required for sustained phosphorylation of tyrosine residues in immunoreceptor tyrosine-based activation motifs (ITAMs) on activating FcγRs (Bakalar et al., 2018). ITAMs are phosphorylated by cytoplasmic membrane-anchored kinase Lyn—a member of the Src family kinases—which remains present within an FcγR-antibody interface due to its lack of extracellular domain (Figure 2e). Lyn is also believed to phosphorylate tyrosines on ITIMs of SIRPα, given previous evidence that Lyn can phosphorylate other ITIM-containing receptors and that depletion of Lyn leads to decreased SIRPα phosphorylation (Abram & Lowell, 2008; Scapini et al., 2009). However, the mechanism by which SIRPα phosphorylation is regulated is not well understood.

We hypothesized that phosphorylation of SIRPα ITIMs, when engaged with CD47, might be similarly regulated by size-dependent exclusion of CD45, since CD45 would not be able to enter the gap to dephosphorylate Lyn-phosphorylated ITIMs. To investigate this, we imaged GPMVs from RAW 264.7 cells on an SLB containing CD47 but not antibody and saw dramatic exclusion of CD45 from interfaces established by only SIRPα-CD47 (Figure 2f). This observation indicates that SIRPα-CD47 binding creates a small enough gap to exclude CD45 and that exclusion of CD45 does not require FcγR engagement.

We next tested whether CD45 exclusion results in increased tyrosine phosphorylation at the interface between macrophages and target particles coated in antibody, CD47, or both ligands (Figure 2g). Using phosphotyrosine immunostaining (not specific to ITAMs or ITIMs), we would expect to detect an increase in phosphorylation at target interfaces with both CD47 and antibody compared to targets with just antibody due to combined labeling of phosphorylated FcγRs ITAMs and SIRPα ITIMs. Surprisingly, phosphorylation at the target particle interface dropped dramatically with the addition of CD47 despite similar levels of antibody on the two target particle populations (Figure 2h). This unexpected result raised the question of why phosphorylation is reduced rather than increased when CD45 is excluded in the presence of CD47.

### SIRPα enrichment decreases phosphorylation at the macrophage-target interface

Our phosphorylation results implied that not only were phosphorylated SIRPα ITIMs not contributing to phosphorylation measured at the particle interface, but that phosphorylation of the antibody-bound FcγR ITAMs was also decreasing. To better quantify this, we immunostained macrophages for phosphotyrosine after allowing them to interact with target particles with increasing antibody:CD47 ratios. Normalizing the average pixel intensity of the phosphotyrosine antibody by the anti-biotin antibody intensity, we find that the level of phosphorylation per anti-biotin antibody at the interface goes down with increasing CD47 (Figure 3a). Importantly, this means that there is proportionally less phosphorylation for each ITAM at the macrophage-target interface. While this correlation suggests that the phosphotyrosine antibody is reporting ITAM phosphorylation, it should be noted that immunostaining may pick up phosphorylation of additional downstream components of the activation pathway. Despite this, the phosphorylation data shows a titratable decrease of activation pathway phosphorylation with increasing CD47.

**Figure 3.**
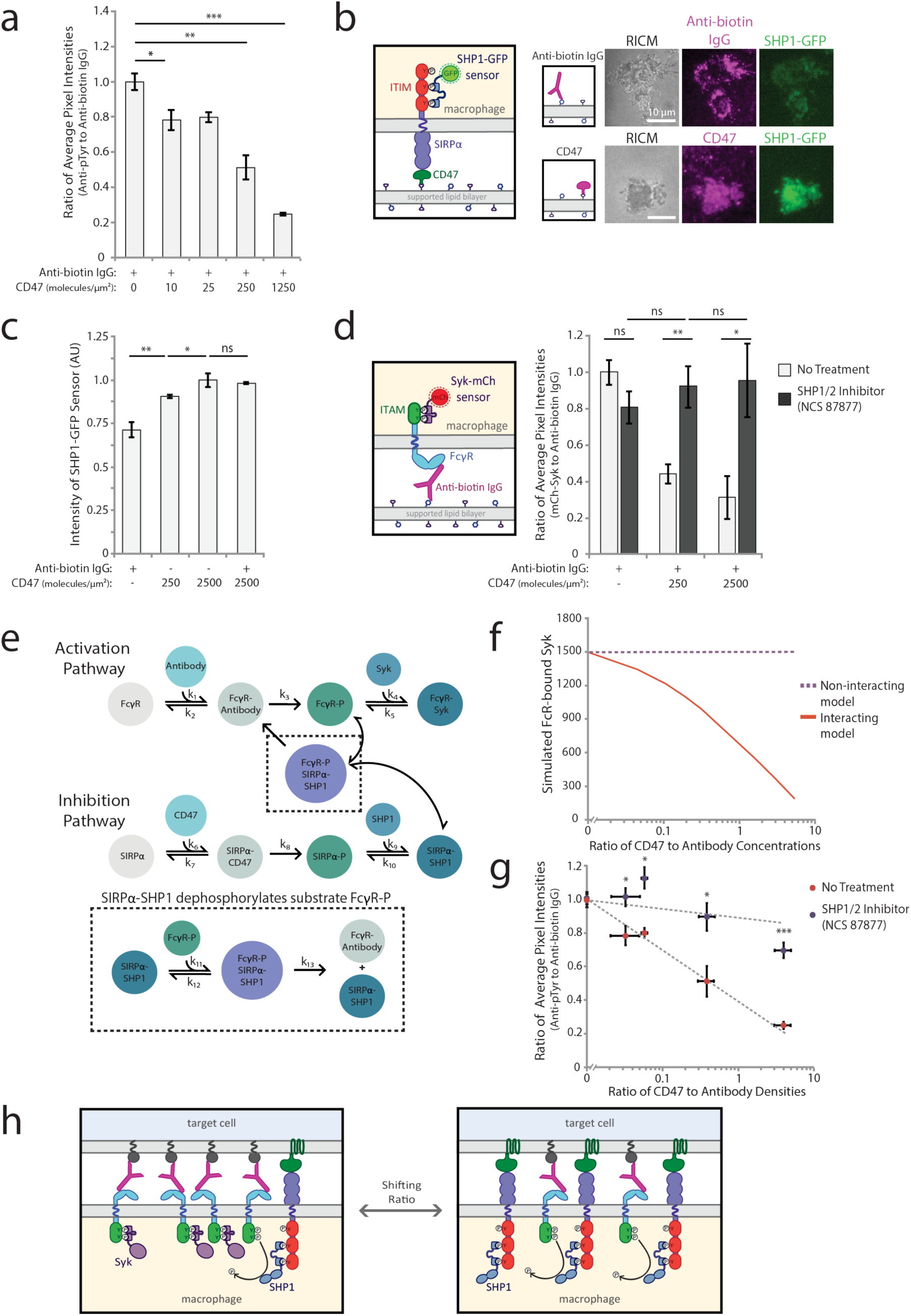
SIRPα-bound SHP1 dephosphorylates activation pathway to inhibit phagocytosis. (a) Quantification of the ratio of anti-phosphotyrosine to anti-biotin IgG fluorescence at macrophage-bead interfaces. RAW 264.7 macrophages interacting target particles coated with a fixed density of AlexaFluor647-labeled anti-biotin IgG (300 molecules/μm^2^) and variable AlexaFluor488-labeled CD47 (0-1250 molecules/μm^2^). Each condition is the average of 3 independent experiments representing a total of >150 particle-cell interfaces. Bars represent mean ± s.e.m. (b) Schematic of SHP1-GFP sensor for detection of phosphorylated SIRPα (left panel). Representative TIRF and RICM images are shown of RAW 264.7 cells expressing SHP1-GFP (green) on SLBs containing either anti-biotin IgG only or CD47 only (each labeled with AlexaFluor647, magenta). (c) Average SHP1-GFP fluorescence signal within cell footprints was quantified for cells on SLBs coated with different CD47 densities (0, 250, 2500 molecules/μm^2^) with and without anti-biotin IgG (300 molecules/μm^2^). Each condition is the average of 3 independent experiments representing a total of >100 macrophage footprints. Bars represent mean ± s.e.m. (d) Schematic of Syk-mCh sensor for detecting phosphorylated FcγRs. RAW 264.7 cells expressing Syk-mCh sensor were imaged on SLBs coated with anti-biotin IgG (300 molecules/μm^2^) and a range of CD47 densities (0, 250, 2500 molecules/μm^2^). Average fluorescence signal within the cell footprint was measured for anti-biotin IgG and Syk-mCh, and the ratio of Syk-mCh to anti-biotin IgG was quantified for each cell. Cell footprint fluorescence was measured for cells with no pre-treatment (light gray bars) and treatment with SHP1/2 phosphatase inhibitor NSC-87877 (dark gray bars). Each condition is the average of 3 experiments representing a total of >200 cell footprints. Bars represent mean ± s.e.m. (e) Schematic of computational model combining activating (top) and inhibitory (bottom) pathways. SIRPα-SHP1 dephosphorylation of FcγR captured in the black dotted inset represents the “interacting” version of the model. (f) “Interacting” and “non-interacting” models were initiated with a fixed antibody concentration and variable CD47 concentrations, yielding a range of CD47:antibody ratios. Each model for each ratio condition was run until steady state was reached, and FcγR-bound Syk was plotted. (g) Ratio of anti-phosphotyrosine to anti-biotin IgG at macrophage-target interfaces was measured for a range of CD47:antibody ratios on target particles (as described in (a)). Quantification was done with and without pre-treatment of macrophages with SHP1/2 phosphatase inhibitor NSC-87877. Each condition is the average of 3 independent experiments representing a total of >150 particle-cell interfaces. Dots represent mean ± s.e.m. (h) Schematic of SHP1-facilitated shutoff of activation pathway. In the absence of CD47, FcγR ITAMs are phosphorylated and bind Syk kinase, promoting phagocytosis. Upon CD47 engagement and SIRPα ITIM phosphorylation, SHP1 phosphatase is brought in close proximity to phosphorylated ITAMs, enabling their dephosphorylation.

To test whether we see a similar result when integrating over the entire macrophage-target interface, we dropped macrophages onto SLBs with and without CD47, and then fixed and immunostained the macrophages for phosphotyrosine (Figure S2a). We again saw proportionally less phosphorylation per antibody enriched in the interface in the condition with CD47. How then does SIRPα-CD47 cause decreased FcγR phosphorylation?

### Close proximity of ITIM-associated phosphatases decreases FcγR phosphorylation

In our immunostained samples, ITAMs and ITIMs are held in close proximity at the interface between a macrophage and target particle due to SIRPα-CD47 and antibody-FcγR co-localizing to similar membrane gaps. We speculated that decreasing phosphorylation in the presence of CD47 did not occur because CD45 was less excluded but rather because soluble phosphatases bind to phosphorylated ITIM domains at the interface (Kharitonenkov et al., 1997; Veillette et al., 1998). If this is the case, both size-dependent exclusion of CD45 and enrichment of ITIMs should affect measured phosphorylation.

Previous work has shown that recruitment of phosphatases SHP1, SHP2, and SHIP to phosphorylated ITIMs is a key regulator of their phosphatase activity (Binstadt et al., 1996; Burshtyn et al., 1997; Neel et al., 2003). SHP1—the most widely studied and directly implicated in down-regulation of macrophage activation—is recruited to phosphorylated SIRPα via its tandem N-terminal SH2 domains and has been shown to be able to dephosphorylate a range of targets, including ITAMs (Frank et al., 2004; Kant et al., 2002). In synthetic peptide studies, SHP1 preferentially dephosphorylates ITAM-like targets of Src kinase, a family member of Lyn kinase, leading to very transient kinase signals in the presence of SHP1 (Frank et al., 2004). In cells, overexpression of SHP1 reduces phosphorylation of activating FcγRIIA and dramatically decreases Syk activation (Huang et al., 2003). Furthermore, following inhibitory stimulus, SHP1 activation peaks approximately 1 min after stimulation and decays in a matter of minutes, even in the sustained presence of the inhibitory stimulus (Huang et al., 2003). This rapid rise and fall of SHP1 activity is consistent with the short timescales of phagocytosis decision-making.

Based on this, we hypothesize that SHP1 dephosphorylates adjacent ITAMs after binding to SIRPα, suppressing FcγR activation. To test whether SHP1 is recruited to the macrophage-target particle interface when CD47 is present, we first created an ITIM phosphorylation sensor by replacing the phosphatase domain of SHP1 with GFP (Figure S2b). This sensor serves to both visualize SHP1 recruitment and also demonstrate SIRPα phosphorylation, which was hard to capture with phosphotyrosine immunostaining. We dropped RAW cells stably expressing this sensor onto SLBs coated in different concentrations of CD47 or antibody (Figure 3b) and quantified the SHP1-GFP sensor intensity within the footprint (Figure 3c). We found that SHP1 sensor recruitment to SLB interfaces containing CD47 was greater than to SLB interfaces with only antibody, with SHP1 recruitment increasing as CD47 surface density increased from 250 to 2500 molecules/μm^2^ and staying approximately constant with the addition of 400 antibody molecules/μm^2^ (Figure 3c). As expected, SHP-1 recruitment decreased upon addition of Src family kinase inhibitor PP2, which should prevent Lyn kinase phosphorylation of ITIMs (Figure S2c). These experiments demonstrate that SHP1 is selectively recruited to SIRPα receptors engaged with CD47 at the target interface, independent of the density of engaged FcγRs.

We next tested if co-localization of ITIMs and ITAMs at the interface between the macrophage and target particle could directly reduce phosphorylation of ITAMs, thereby shutting off FcγR activation. Syk kinase is recruited to phosphorylated FcγR ITAMs via its two SH2 domains and amplifies the activation signaling cascade leading to phagocytosis by phosphorylating numerous downstream targets (Crowley et al., 1997; Fütterer et al., 1998; Tsang et al., 2008). However, the presence of SHP1 on nearby ITIMs could dephosphorylate ITAMs, blocking Syk recruitment and downstream signaling. We have already demonstrated that bound SIRPα-CD47 physically fit into the same membrane gap created by FcγR-antibody binding, allowing activating and inhibitory receptors to intermingle (Figure 2d), and we have also shown that CD45 is excluded from that gap, allowing both ITIMs and ITAMs to be phosphorylated by Lyn kinase (Figure 2f).

To test if local Syk recruitment decreases as a result of increasing CD47, we added macrophages to SLBs with different antibody:CD47 ratios and measured the signal from a fluorescent Syk sensor, which we previously developed to monitor phosphorylated ITAMs in real time (Bakalar et al., 2018) (Figure 3d, left panel). Adding 250 molecules/μm^2^ of CD47 to the bilayer dramatically decreased the amount of Syk recruited, and adding 2500 molecules/μm^2^ decreased recruitment even further (Figure 3d, right panel, light bars). This data shows that in the presence of CD47, the same amount of antibody yields markedly less Syk binding, suggesting direct dephosphorylation of ITAMs by SHP1.

### Model of ITIM-ITAM negative feedback captures ratio-dependent FcγR phosphorylation

With experimental evidence of negative feedback resulting from CD47 enrichment at the macrophage-target particle interface, we wondered whether a model of SHP1 recruitment leading to decreased FcγR phosphorylation and subsequent Syk recruitment could capture the non-linear behavior of ratio-dependent phagocytosis (as observed in Figure 1d). To investigate this, we created a simple kinetic model that includes the parallel pathways of activation and inhibition (Figure 3e, Table S1). Each pathway is based on receptor-ligand binding, which drives receptor phosphorylation and subsequent binding of downstream effectors, SHP1 or Syk. Our model assumes that binding drives receptor enrichment and CD45 exclusion, enabling receptor phosphorylation.

We tested two versions of the model, one in which activation and inhibition are interacting and one in which they are not. In the ‘non-interacting’ version, the pathways proceed independently, with no SHP1 negative feedback on FcγR ITAM phosphorylation. In the ‘interacting’ version, activation and inhibition pathways are connected by a single integration point, dephosphorylation of FcγR ITAMs by SIRPα-bound SHP1 (Figure 3e). Constants used in the model were obtained from the literature, when available, and others were estimated (Table S2) (Barua et al., 2012; Brooke et al., 2004; Li et al., 2007; Ren et al., 2011; Selner et al., 2014). The model is initiated with defined concentrations of unbound CD47, antibody, SIRPα, FcγR, SHP1, and Syk. FcγR-bound Syk is used as a proxy for phagocytosis since its localization to the target interface is known to be required for activation pathway signaling (Crowley et al., 1997; Kiefer et al., 1998). As expected, the ‘non-interacting’ model shows that FcγR-bound Syk is insensitive to varying ratio of CD47:antibody concentrations (Figure 3f). However, for the ‘interacting’ model, SHP1 is able to dephosphorylate FcγR ITAMs and FcγR-bound Syk decreases sharply with increasing CD47:antibody ratio.

To experimentally test the ‘non-interacting’ model in cells, we incubated macrophages with target particles coated in a range of CD47:antibody ratios (as in Figure 3a) either with or without SHP1/2 inhibitor NSC-87877, which has previously been shown to be specific (Chen et al., 2006). Addition of the inhibitor should prevent SHP1 from dephosphorylating FcγRs and other activation pathway components, allowing phosphorylation to remain high at the target interface. Consistent with our ‘non-interacting’ model, we found that the amount of phosphorylation per antibody is higher in the presence of inhibitor (versus the control) for each CD47:antibody ratio (Figure 3g), highlighting the key role that SIRPα-bound phosphatases play in shutting down phagocytosis. To test the effect of SHP1/2 inhibition on Syk recruitment, we dropped cells onto SLBs with varying CD47:antibody ratios in the absence or presence of SHP1/2 inhibitor NSC-87877. Syk sensor recruitment in the presence of the inhibitor stayed constant for conditions including or excluding CD47, as expected from the ‘non-interacting’ model (Figure 3d, right panel, dark bars).

Both the ITIM phosphorylation and Syk recruitment data point to a model of macrophage decision-making in which small changes in CD47:antibody ratio drive shifts in relative enrichment of SIRPα and FcγRs (Figure 3h). This changing SIRPα enrichment alters the relative density of ITIM-bound phosphatases, which can then rapidly and effectively dephosphorylate nearby ITAMs to stop phagocytosis.

### ITAM:ITIM ratio dictates phagocytosis levels independent of extracellular identities

The critical parameter in our model is the number of antibody-bound FcγRs relative to the number of CD47-bound SIRPα, which we can abstract to mean the ratio of ITAM:ITIM enriched at a target interface. Hence, we wondered whether ratio-dependent phagocytosis was specific to the competition between FcγRs and SIRPα or whether any receptor with an ITAM or ITIM would behave similarly. We hypothesized that artificially shifting the ITAM:ITIM ratio at the target interface should alter phagocytosis independent of the extracellular binding domain identities that drive receptor enrichment.

To test this, we created a chimeric receptor containing the intracellular ITAM-containing portion of FcγRIIA (Figure S3a). As the extracellular binder, we used the Syn18 leucine zipper sequence from the SynZip toolkit, which offers an orthogonal means of binding a target (Thompson et al., 2012). As a ligand, we used the complementary SynZip motif, Syn17, attached to three repeats of FNIII domain (Fibcon), which we used previously as a tool for modulating extracellular protein height (Bakalar et al., 2018), and a His-tag for attachment to the SLB (Syn17-F3L). Upon binding, this interaction should achieve a membrane gap similar to that of SIRPα-CD47, approximately 13 nm. Using target particles coated in Syn17-F3L, we found that these receptors could drive efficient phagocytosis and that phagocytosis was specific to the FcγRIIA ITAM motif (Figure S3b). When CD47 and Syn17-F3L were added in varying ratios on target particles, we found that the synthetic FcγRIIA receptor responded in a titratable way to the activation:inhibition ratio, very similar to the endogenous receptors (Figure 4a).

**Figure 4.**
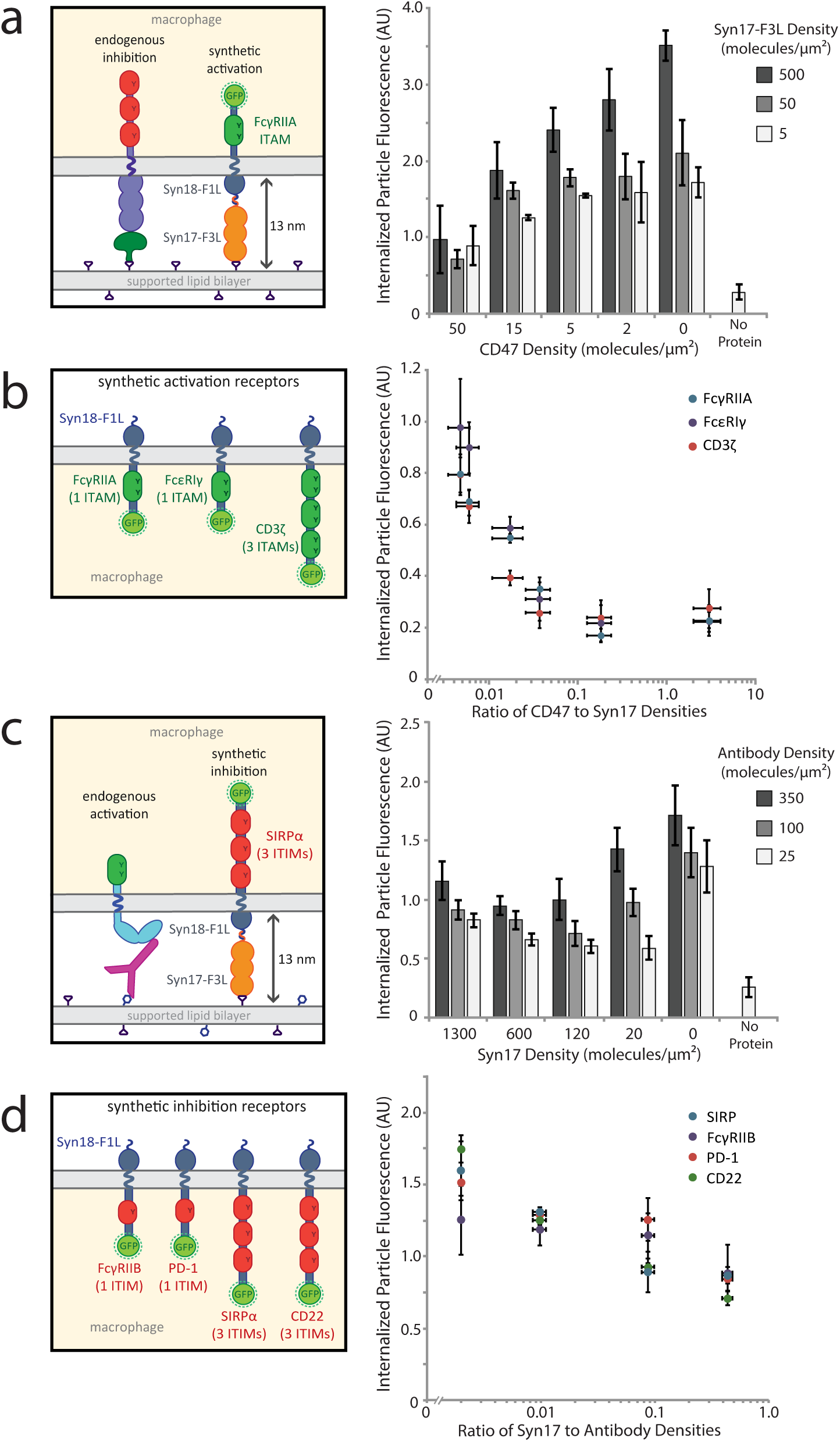
Ratio of activation to inhibition governs phagocytosis in a synthetic context. (a) Target particles coated in varying ratios of Syn17-F3L and CD47 were incubated with RAW 264.7 macrophages stably expressing the synthetic activating receptor Syn18-F1L-TM-FcγRIIA-GFP. For each density of Syn17-F3L (5, 50, 500 molecules/μm^2^), five densities of CD47 (0-50 molecules/μm^2^) were added. Each condition is the average of three independent experiments representing a total of >300 cells. Bars represent mean ± s.e.m. (b) RAW 264.7 macrophages stably expressing synthetic receptors containing three different ITAM-containing signaling motifs: FcγRIIA, CD3ζ, and FcεRI γ-chain. Phagocytosis was quantified for target particles with a range of CD47:Syn17-F3L density ratios. Each condition is the average of three independent experiments representing a total of >300 cells. Dots represent mean ± s.e.m. (c) Target particles containing varying ratios of Syn17-F3L and anti-biotin IgG were incubated with RAW 264.7 macrophages stably expressing synthetic inhibitory receptor Syn18-F1L-TM-SIRPα-GFP. In contrast to (A) and (B), note that Syn17-F3L is now acting as an inhibitory ligand. For each density of anti-biotin IgG added (25, 100, 350 molecules/μm^2^), five densities of Syn17-F3L (0-1300 molecules/μm^2^) were added. Each condition is the average of three independent experiments representing a total of >300 cells. Bars represent mean ± s.e.m. (d) RAW 264.7 macrophages stably expressing synthetic receptors containing four different ITIM-containing signaling motifs: FcγRIIB, SIRPα, PD-1, and CD22 Phagocytosis was quantified for target particles with a range of Syn17-F3L:anti-biotin IgG density ratios. Each condition is the average of three independent experiments representing a total of >300 cells. Dots represent mean ± s.e.m.

To investigate whether this ratio-dependent signaling was specific to the FcγRIIA ITAM domain or whether ITAMs from other receptors would have the same effect, we created additional chimeric activating receptors featuring the intracellular signaling regions of the CD3 ζ-chain of the T-cell receptor complex and FcεRI γ-chain (Figure S3a). Each receptor construct was identical, with the exception of the signaling domain. We found that macrophages expressing each chimeric receptor responded similarly to Syn17-3L alone (Figure S3c) and that each receptor responded similarly to a titration of CD47, suggesting that ratio dominates phagocytosis independent of the receptor identities (Figure 4b). Interestingly, the ratio of Syn17 to CD47 required to get phagocytosis over background was similar to that of antibody to CD47, approximately 10:1, indicating that ratiometric signaling through synthetic receptors mimics that through endogenous FcγRs.

To determine if similar rules exist for ITIM-containing domains, we next created a chimeric SIRPα receptor to compete against the activation signal of endogenous FcγRs. The receptor design was identical to the activation chimeric receptors, except for replacing ITAM-containing domains with SIRPα’s ITIM-containing intracellular region (Figure S3a). We found that the chimeric SIRPα response is also titratable with ratio similar to the endogenous SIRPα (Figure 4c). Additional inhibitory chimeric receptors featuring the ITIMs of CD22, FcγRIIB, and PD-1 also showed ratio-dependent phagocytosis behavior (Figure 4d). Interestingly, we observed that macrophage phagocytosis with the chimeric SIRPα never fully decreased to zero in response to the now inhibitory Syn17-F3L ligand. We attribute this elevated background engulfment to the high affinity and enrichment of Syn17-F3L to the Syn18 receptor, independent of the inhibitory signaling (Figure S3e). Taken together, these experiments with chimeric receptors demonstrate that ratio-dependent phagocytic signaling is generalizable to other ITAM and ITIM signaling domains.

### Ratio-dependent phagocytosis of tumor cells can be shifted using blocking antibodies

Finally, we tested whether our model of ratio-dependent signaling was applicable to live tumor cells and could guide use of targeting and blocking antibodies to enhance phagocytosis. Tumor cell surfaces are significantly more complex than the two-component reconstituted target particles used in our assays above and therefore may be subject to additional inhibitory interactions that further complicate efforts to overcome inhibitory signaling (Feng et al., 2019).

To conduct tumor cell phagocytosis assays, we used mouse bone marrow derived macrophages (BMDMs) and the mouse colon adenocarcinoma cell line MC38, both of which have been used in previously published tumor cell eating experiments (Morrissey & Vale, 2019; Xu et al., 2017). MC38 cells were stably expressing human HER2 protein (MC38-hHER2), a well-characterized tumor antigen over-expressed in 15-30% of breast cancers (Iqbal & Iqbal, 2014). We opsonized the MC38-hHER2 cells with a mouse anti-hHER2 antibody, which was modified from the clinical hHER2-targeting antibody Trastuzumab (McKeage & Perry, 2002) and a kind gift from Aduro Biotech. The opsonized tumor cells were incubated with BMDMs for 30 minutes before imaging (Figure 5a). Importantly, we estimate that macrophage FcγR and SIRPα receptors are in excess of their corresponding ligands on MC38 cells.

**Figure 5.**
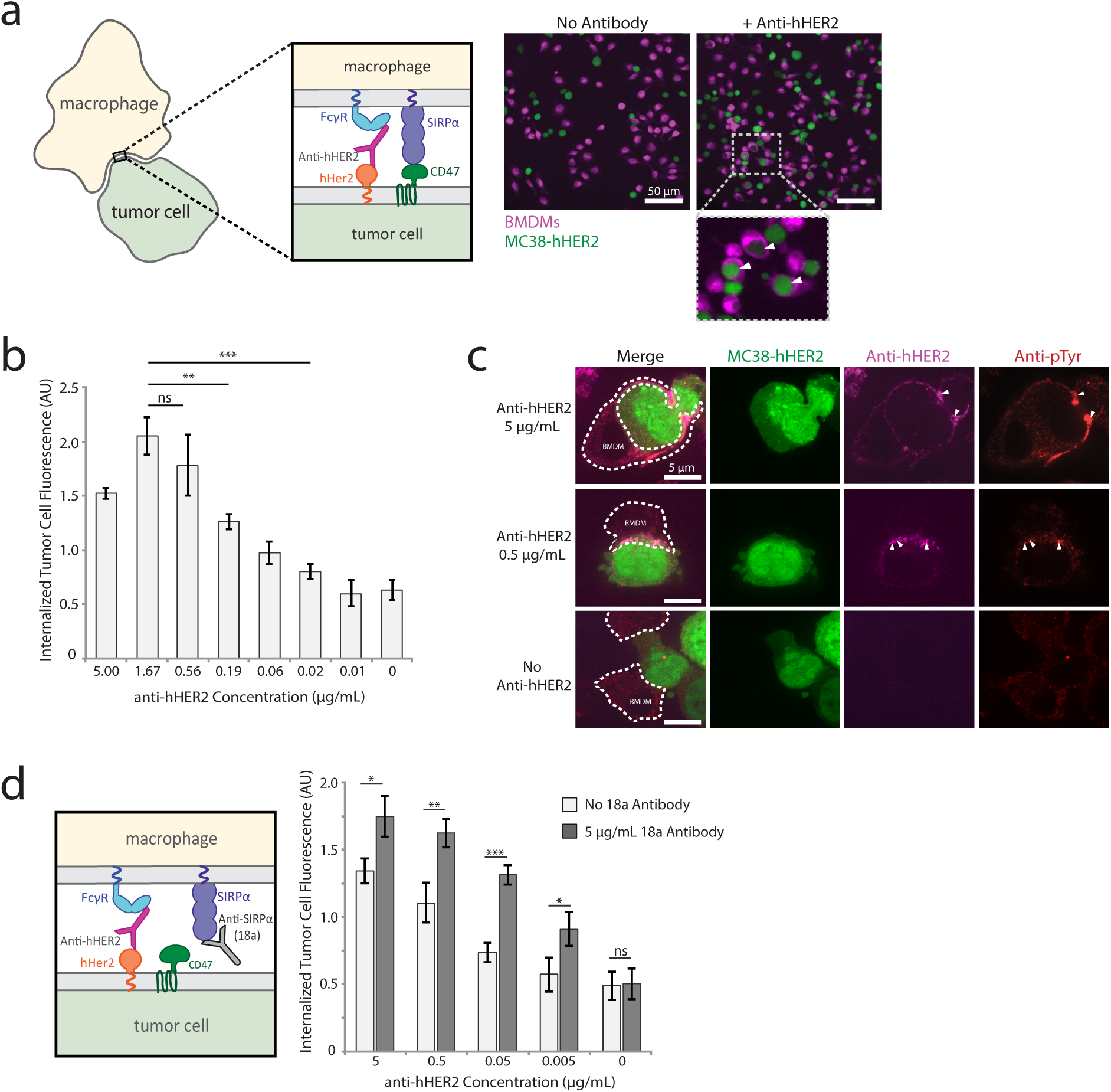
Tumor cell phagocytosis is dictated by activation-inhibition ratio. (a) Bone marrow derived macrophages (BMDMs) were incubated with MC38-hHER2 cells either with or without tumor targeting anti-hHER2 antibody opsonization. Representative confocal images are shown. BMDMs were labeled with Cell Tracker Deep Red (magenta) and MC38-hHER2 cells were labeled with CellTracker Green (green). White arrows within inset indicate phagocytosed MC38-hHER2 cells. (b) MC38-hHER2 cells were opsonized with a range of anti-hHER2 antibody concentrations (0-5 μg/mL) and incubated with BMDMs. Average tumor cell phagocytosis was quantified. Each condition is the average of 3 independent experiments representing >150 total cells. Bars represent mean ± s.e.m. (c) Fluorescent confocal images of fixed BMDMs engaging with MC38-hHER2 cells opsonized with different concentrations of anti-hHER2 antibody (0, 0.5, 5 μg/mL). MC38-hHER2 cells were labeled with CellTracker Green (green), anti-hHER2 with AlexaFluor647 (magenta) and phosphorylation with anti-phosphotyrosine (visualized through AlexaFluor546-labeled secondary antibody, red). BMDM outlines are shown in white dotted lines. Areas of high anti-hHER2 enrichment are highlighted with white arrows. (d) Schematic of anti-SIRPα blocking antibody 18a interrupting SIRPα binding to CD47 (left panel). MC38-hHER2 cells incubated with a range of anti-hHER2 antibody concentrations (0-5 μg/mL) were added to BMDMs in the presence or absence of 5 μg/mL 18a antibody. Average tumor cell phagocytosis was quantified. Each condition is the average of 3 independent experiments representing >150 total cells. Bars represent mean ± s.e.m.

In order to shift the activation:inhibition ratio we chose to vary the activation signal by titrating the anti-hHER2 antibody, while CD47-SIRP interaction stayed constant. This strategy was informed by clinical evidence that efficacy of tumor growth inhibition is correlated to HER2 opsonization levels (McLarty et al., 2009). Over an anti-hHER2 concentration range of 0.01 - 5 μg/mL, we see a clear increase in tumor cell phagocytosis, consistent with an increased activation:inhibition ratio (Figure 5b). The highest concentration (5 μg/mL) showed a slight decrease in phagocytosis, potentially due to excess anti-hHER2 antibody in solution saturating unbound FcγRs and preventing their engagement with anti-hHER2 on the tumor cell. To confirm that increased opsonization is accompanied by changes in receptor phosphorylation, we fixed BMDMs interacting with MC38-hHER2 cells and immunostained them for phosphotyrosine. Consistent with our reconstituted target particle experiments, we find that phosphorylation is concentrated at the interface between the BMDM and tumor cell, and that regions of antibody enrichment correlate with high levels of phosphorylation (Figure 5c).

Both BMDMs and RAW 264.7 cells exhibit a similar sensitivity to antibody:CD47 ratios on reconstituted target particles (Figure S4a), consistent with BMDMs and RAW cells having comparable ratios of FcγR:SIRPα, roughly 2:3 and 1:2, respectively, despite having different absolute numbers of receptors (Figure S4b). From surface measurements of MC38-hHER2 cells, we estimated 26,100 hHER2 molecules/cell and 30,100 CD47 molecules per cell. Surprisingly, at antibody saturation of hHER2, this yields an activation:inhibition ratio of approximately 1:1, much lower than the 10:1 required for our target particles. This difference in ratio may be due to a number of reasons, including differences in antibody affinity and partial segregation of SIRPα-CD47 from antibody-bound FcγR resulting from the size of HER2, suggesting that the threshold ratio must be determined for each specific antibody and antigen.

Finally, we asked if we could shift the activation-inhibition ratio to improve phagocytic efficiency at lower anti-hHER2 antibody concentrations by blocking SIRPα-CD47 interactions and tipping the balance towards activation. This can be done most directly with antibodies that either block CD47 or SIRPα binding. Since CD47 is present on all cells (including macrophages) and high concentration of a CD47 blocking antibody could have side effects (Russ et al., 2018), we chose to target SIRPα. CD47 blocking antibodies also have the potential to interact with macrophage FcγRs via their Fc tails and thereby complicate interpretation of phagocytosis results, so targeting SIRPα over CD47 provided an experimental advantage as well. Previously published SIRPα blocking antibodies, such as the widely-used p84 clone, have been shown to have only partial or limited blocking properties (Yanagita et al., 2017). Since no anti-mouse SIRPα blocking antibodies are currently commercially available to our knowledge, we utilized a new SIRPα antibody, 18a, developed by Aduro Biotech to bind mouse SIRPα and block CD47 engagement. We found that adding 5 μg/mL of 18a antibody over multiple dilutions of anti-hHER2 antibody systematically raised phagocytosis levels for each concentration of anti-hHER2 (Figure 5d), indicating that phagocytosis of tumor cells is, like the reconstituted target particles, dependent on antibody:CD47 ratios. This data also suggests that the SIRPα-CD47 axis is a primary means of inhibition for these tumor cells, since ratiometrically blocking this interaction increases phagocytosis significantly.

## DISCUSSION

Immune cells, including macrophages, make important effector function decisions by directly interacting with target cells. Receptor engagement, enrichment, and signaling at cell-cell contacts are critical for determining how an immune cell responds to a target, and understanding these steps at a mechanistic level has the potential to inform efforts to harness macrophage effector functions for therapeutic purposes. This is especially the case when there is competition between inhibitory and activating cues, where the immune cell must make a decision based on inputs from receptor signaling. Here, we ask how macrophages process activating signals from antibodies and inhibitory signals from CD47 on target particles. We show quantitatively that the activating and inhibitory signals produced by receptor enrichment at the macrophage phagocytic synapse are integrated in a ratio-dependent manner, with approximately 10 antigen-bound antibodies required to overcome one CD47. Rather than using a threshold number of activating or inhibitory molecules to make a go/no-go decision, macrophages use the relative number of ITAMs and ITIMs enriched at an interface. Since each antibody-bound activating FcγR brings one or two ITAMs to the synapse while each CD47-bound SIRPα brings three ITIMs, our findings show that phagocytosis proceeds when there are at least 2-3 ITAMs for every ITIM.

The majority of targets encountered by a macrophage are likely to present “obvious” choices in which one signaling pathway overwhelmingly dominates. In the event of heavily antibody-opsonized pathogens, for example, there is likely little engagement of inhibitory checkpoints and high levels of activating signal. Conversely, most healthy cells in our bodies have few activating ligands but high levels of CD47 and other inhibitory checkpoint ligands. However, diseases of self, such as cancer and autoimmune diseases, present an interesting and complex gray area, where tumor antigen opsonization must compete with CD47, leaving immune cells to process high levels of both signals. This study examines ligand densities across a regime in which macrophages transition between inhibition and activation in order to obtain a mechanistic picture of how macrophages respond to complex stimuli.

We find that the ratio of antibody to CD47 dominates phagocytosis decisions in ADCP and that phosphatase feedback is a key reason for this. Ultimately, competition between activation and inhibition comes down to a race between phosphorylation and dephosphorylation, with Syk and SHP1 in direct competition. Subtle changes in ligand ratio drive changes in relative recruitment of SHP1 (and other phosphatases such as SHP2 and SHIP) to ITIMs, which rapidly shuts down activation by dephosphorylating nearby ITAMs. Importantly, SHP1 likely dephosphorylates ITIMs as well, as evidenced by low phosphorylation measurements at the interface between macrophages and targets, even in the presence of high SIRPα-CD47 engagement. This suggests that the phosphatase feedback acts not only to shut down activation but also to quickly turn off inhibition, returning the cell to a neutral state, poised to respond to the next potential target. The impact of ratio is amplified by SHP1’s efficiency as an enzyme: the *k_cat_* of SHP1 is as much as 60 times greater than that of maximally activated Syk (Barua et al., 2012; Selner et al., 2014; Tsang et al., 2008). A similar negative feedback model was proposed for PD-1, in which SHP2 phosphatase recruited to phosphorylated PD-1 clusters suppresses T cell receptor activation (Yokosuka et al., 2012). As such, we speculate that trans-dephosphorylation of activating pathways by proximal phosphatases may be a shut-off mechanism universal to all ITIM motifs.

Interestingly, it appears that the SIRPa-CD47 inhibitory checkpoint recruits SHP1 when CD45 is excluded from the interface, presumably because the gap formed drives size-dependent exclusion of CD45 as originally proposed by Davis and van der Merwe for T cell receptor activation (Davis & van der Merwe, 2006). Our previous work showed that antigen size is critical for effective ADCP, with shorter antigens more efficiently excluding CD45, which in turn drives increased ITAM phosphorylation (Bakalar et al., 2018). We speculate that the similarity in SIRPα-CD47 height for engagement and FcγR-antibody-antigen height for efficient phagocytosis is not coincidental but rather has evolved specifically to shut down phagocytosis at its most efficient point if markers of self are detected. Our findings point to the possibility that CD45’s main role is to maintain non-interacting macrophages in an “off” state by preventing both activating and inhibitory signaling. Exclusion of CD45 upon close contact with a target then enables engaged and enriched receptors of either pathway to signal, with FcγR and SIRPα signaling interpreted ratiometrically.

In our model of macrophage FcγR decision-making, co-localization of SIRPα-CD47 and FcγR-antibody binders is not only convenient for CD45 exclusion but also important to provide SHP1 access to phosphorylated ITAMs. If the two species were spatially segregated but still excluding CD45, we might expect SIRPα-CD47 to be a less potent inhibitor since SIRPα-bound SHP1 would have less access to phosphorylated ITAMs. Consistent with this idea, a study in NK cells found that inhibition became less potent when the inhibitory KIR2DL1-HLAC interaction was made taller (Köhler et al., 2010), potentially because this caused KIR2DL1-HLAC to be segregated from activating NKG2D-MICA interactions at the interface. The importance of co-localization is also seen in our SLB measurements of macrophage footprints in the presence of increasing CD47 densities. Despite a similar average enrichment of antibody within a footprint, we see a decrease in antibody clustering and an overall decline in phosphorylation. This implies that local organization of the two binders is shifted as the ratio changes, which may facilitate inhibitory shut-down of activation through dephosphorylation.

Our data also suggests that the most important node for integrating activating versus inhibitory signaling is at the receptors themselves. In NK cells, it has been suggested that Vav1 acts as an integrative node for inhibition triggered by inhibitory KIR receptors (Stebbins et al., 2003). However, macrophages lacking Vav1 are still able to phagocytose (Hall et al., 2006). In fact, many commonly cited players in downstream phagocytosis signaling can be removed without preventing productive FcγR-mediated phagocytosis, including WAVE and Rho proteins (Caron & Hall, 1998; Kheir et al., 2005). This suggests that phagocytosis, which makes use of a complex activation network, is not dependent on a single integrator downstream of the receptors. Furthermore, phagocytosis decisions are made very quickly, and target particles in our assays are frequently phagocytosed in a matter of seconds after contact, a timeframe consistent with the rapid dephosphorylation of FcγR ITAMs by phosphatases recruited to SIRPα ITIMs needed to block phagocytosis. This dephosphorylation may be further modulated by downstream processes that have been previously associated with CD47 on targets, including myosin phosphorylation and integrin activation (Morrissey & Vale, 2019; Tsai & Discher, 2008).

To show the relevance of our ratiometric model beyond reconstituted target particles, we demonstrated that live tumor cell phagocytosis follows the same requirement for activating:inhibitory signaling. Though the tumor cells’ have a far more complex cell surface, our study indicates that careful characterization of receptor and ligand densities on both immune and tumor cells may be important considerations for immunotherapies. More specifically, designing individualized therapeutic dosing informed by surface densities of tumor antigen, CD47, SIRPα, and FcγR could help to optimize results for specific cancers that involve macrophage effector function. The results of our study emphasize that small changes in antibody:CD47 ratio can have important effects on phagocytosis efficiency, something that may be important to consider as part of rationally designed combinatorial checkpoint blockade treatments.

Interestingly, we saw that the ratio of activating:inhibitory ligand necessary for phagocytosis was much lower in tumor cell experiments than in our reconstituted system, approximately a 1:1 ratio versus a 10:1 ratio, respectively. This difference could be due to variation in antibody affinity towards the antigen or the FcγR, leading to different enrichment profiles and potentially higher activation. CD47 may also be less accessible for SIRPα in the crowded environment of tumor cell membranes than in our reconstituted target particle experiments, leading to lower inhibition. In addition, hHER2 is estimated to stand 11 nm above the cell membrane (Bakalar et al., 2018), meaning hHER2-antibody-FcγR would be expected to have a greater height than bound CD47-SIRPα. This could potentially lead to size-dependent spatial separation of the activating and inhibitory signals, which could reduce the ability of SHP1 recruited to ITIMs to dephosphorylate adjacent ITAMs, as we speculate above. The complexity of the tumor cell surface may also lead to a more stable macrophage-target interface facilitated by additional receptor-ligand interactions not present in the reconstituted system. Though T-cell immunological synapse adhesion and stability has been extensively studied (Dustin, 2007; Graf et al., 2007; Philipsen et al., 2013), the impact of macrophage interface stability on signaling efficiency has not been widely characterized and presents a topic for further study.

The ratiometric signaling model we propose here may be broadly relevant to ITAM-dependent signaling in different cell types. In previous work in NK cells, overexpression of inhibitory Ly49 ligands on model tumor cells was able to shut down NKG2D-mediated activation in an expression-dependent manner, suggesting that relative amounts of the activating and inhibitory ligands were important in that system (Malarkannan, 2006). Additionally, previous work in microglia has shown that complement receptor subunit CD11b requires ITAM-containing adapter protein DAP12 (Wakselman et al., 2008), indicating that not only could ITAM:ITIM ratio be important in complement-mediated phagocytosis but potentially relevant to non-immunoreceptor (e.g., integrin) signaling as well.

However, whether ratio-dependent competition of activating and inhibitory signaling is widely applicable to diverse immune cell types will require careful quantification of effector function over a range of absolute receptor surface densities. Furthermore, while our *in vitro* studies provide insight into mechanism that can influence receptor spatial organization, how this clustering affects immunological synapses *in vivo* requires further investigation. Though extracellular binding domains of immune receptors have been honed for target specificity and affinity, and ITAM or ITIM sequence variability might alter downstream binding affinities, this study leads us to speculate that diverse immune decision-making acts through a common mechanism that involves receptor binding, enriching, and subsequently signaling in a ratio-dependent manner.

## MATERIALS AND METHODS

### Macrophage and tumor cell culture

RAW 264.7 macrophage-like cell line was obtained from the UC Berkeley Cell Culture Facility. Cells were cultured in RPMI 1640 media (Corning) supplemented with 10% heat-inactivated fetal bovine serum (HI-FBS, Thermo Fisher Scientific) and 1% Pen-Strep (Thermo Fisher Scientific) in non-tissue culture-treated 10 cm dishes (VWR) at 37°C, 5% CO_2_.

Bone marrow derived macrophages (BMDMs) from C57BL/6 (B6) mice were a kind gift from the Portnoy Lab (UC Berkeley). BMDMs were grown in RPMI 1640 media supplemented with 10% HI-FBS and 1% Pen-Strep at 37°C. BMDMs were used in experiments within 24 hours of thawing.

MC38 cells stably expressing human HER2 (referred to as MC38-hHER2) were obtained from Aduro Biotech. MC38-hHER2 cells were grown in RPMI 1640 media supplemented with 10% HI-FBS and 1% Pen-Strep at 37°C.

HEK 293T cells were obtained from the Cell Culture facility at UC San Francisco. Cells were cultured in DMEM media (Thermo Fisher Scientific) supplemented with 10% HI-FBS and 1% Pen-Strep at 37°C.

All cell lines tested negative for mycoplasma as verified by Mycoalert detection kit (Lonza).

### Preparation of cell-like reconstituted target particles

Cell-like reconstituted target-particles were generated according to previously published protocol (Joffe, A. et al., 2020), summarized in the following sections.

#### Formation of small unilamellar vesicles (SUVs)

SUVs were prepared by rehydrating a lipid film composed primarily of POPC (1-palmitoyl-2-oleoyl-sn-glycero-3-phosphocholine, Avanti Polar Lipids), doped with 0.5% of DPPE-biotin (1,2-dipalmitoyl-sn-glycero-3-phosphoethanolamine-N-(biotinyl), Avanti Polar Lipids), 1% DGS-Ni-NTA (1,2-dioleoyl-sn-glycero-3-[(N-(5-amino1-carboxypentyl)iminodiacetic acid)succinyl] with nickel salt, Avanti Polar Lipids), and 0.4% LISS-Rhodamine (1,2-dioleoyl-sn-glycero-3-phosphoethanolamine-N-(lissamine rhodamine B sulfonyl), Avanti Polar Lipids) in pure deionized H_2_O. The rehydrated solution was vortexed briefly and then sonicated using a tip-sonicator at 20% power pulsing off and on for 3 minutes. SUVs were then filtered through a 200 nm PTFE filter (Millipore). SUV solutions were stored at 4°C and used within 48 hours of preparation to avoid phospholipid oxidization.

#### Formation of cell-like target particles

40 μL of 4.07 μm silica beads (Bangs Laboratories) were cleaned using a 3:2 mixture of H_2_SO_4_:H_2_O_2_ (Piranha) for 30 minutes while sonicating. Cleaned beads were spun down at 1000 x g and washed 3 times with 1 mL pure water before being resuspended in 400 μL of water. Cleaned beads were stored at room temperature. To form SLBs, 40 μL of SUVs was added to 160 μL of MOPS buffer (25 mM MOPS (3-(N-morpholino)propanesulfonic acid), Thermo Fisher Scientific), 125 mM NaCl, pH 7.4), along with 20 μL of clean bead slurry. The mixture was incubated at room temperature for 15 minutes while rotating continuously. Coated beads (also referred to as target particles) were spun down at 50 x g for 1 minute, and 200 μL of the solution was removed and replaced with PBS (Corning).

#### Addition and quantification of protein on target particles

AlexaFluor647-labeled anti-biotin mouse IgG (clone BK-1/39, Thermo Fisher Scientific) and AlexaFluor488-labeled His-tagged recombinant mouse CD47 (SinoBiological) were diluted in PBS to appropriate concentrations for target particle experiments. 50 μL of SLB-coated beads was added to 50μL of protein dilution, and beads were incubated at room temperature for 20 minutes with continuous rotation.

SLB and protein-coated target particles were diluted in PBS and analyzed using an Attune NxT flow cytometer (Thermo Fisher Scientific). Target particles were compared to calibrated beads with known numbers of AlexaFluor488 and AlexaFluor647 fluorophores (Quantum MESF Kits, Bangs Laboratories), which enabled calculate of protein surface density.

#### Fluorescent labeling of proteins

His-tagged recombinant mouse CD47 (as well as proteins and antibodies noted later) was labeled using AlexaFluor488 NHS Ester (Succinimidyl Ester, Thermo Fisher Scientific) reconstituted in anhydrous DMSO (dimethylsulfoxide, Sigma Aldrich). Dye was mixed with protein at a 5x molar ratio (dye:protein ratio was 5:1) and incubated at room temperature for 1 hour. Excess dye was removed by purifying protein over NAP-5 columns (GE Healthcare). Labeling was confirmed using NanoDrop 2000c (Thermo Fisher).

### Imaging techniques

All live cells were maintained at 37°C, 5% CO2 with a stage top incubator (Okolab) during imaging. For confocal microscopy, cells were imaged with a spinning disk confocal microscope (Eclipse Ti, Nikon) with a spinning disk (Yokogawa CSU-X, Andor), CMOS camera (Zyla, Andor), and either a 20x objective (Plano Fluor, 0.45NA, Nikonor a 60x objective (Apo TIRF, 1.49NA, oil, Nikon). For total internal reflection fluorescence (TIRF) microscopy, cells were imaged with TIRF microscope (Eclipse Ti, Nikon), 60x objective (Apo TIRF, 1.49NA, oil, Nikon) and EMCCD camera (iXON Ultra, Andor). Both microscopes were controlled with Micro-Manager. Images were analyzed and prepared using FIJI (Schindelin, J. et al., 2012).

### Phagocytosis assays

#### Target particle phagocytosis assay

35,000 macrophages were seeded in wells of a tissue-culture flat-bottom 96-well plate (Falcon) in 100 μL of RPMI 1640 medium. Post-seeding, cells were incubated at 37°C for 3-4 hours prior to target particle addition. 100 μL of target particles were prepared for each well with appropriate protein concentrations, and 95 μL of the 100 μL were added. Macrophages were incubated with target particles at 37°C for exactly 20 minutes. During those 20 minutes, the remaining 5μL were diluted in 250 μL of PBS and immediately measured using flow cytometry as previously described.

After incubation, wells were washed twice with PBS to remove excess particles. PBS containing 1 μM of CellTracker Green (CMFDA, Thermo Fisher Scientific) and 10 μM Hoechst 33342 (Thermo Fisher Scientific) was added to wells to stain the cytoplasm and nuclei, respectively. Wells were imaged after 10 minutes of staining using a spinning-disk confocal microscope (Nikon) at 20x. Images were acquired in an automated grid pattern at the same location within each well to reduce bias in image acquisition. For each well, at least 100 cells were imaged. Images were than analyzed using a custom CellProfiler (v2.1.1, Broad Institute (Carpenter, A.E. et al., 2006)) project program to identify single cells and quantify the internalized target particle fluorescence intensity (of LISS-Rhodamine lipid) within each cell. Average internalized fluorescence per cell was calculated per condition. Independent replicates were conducted on different days, and replicates were normalized to average eating for all conditions to control for day-to-day variation.

#### Tumor cell phagocytosis assay

After thawing and resuspending in RPMI 1640 medium, BMDMs were incubated with 1 μM of CellTracker Deep Red (Thermo Fisher Scientific) at room temperature for 10 minutes. Cells were washed 2 times in PBS to remove excess dye, resuspended in RPMI 1640 medium and seeded at 40,000 cells per well in a tissue-culture flat-bottom 96-well plate. Cells were incubated at 37°C overnight (approximately 18 hours prior to tumor cell addition).

MC38-hHER2 cells were incubated in 1 μM Cell Tracker Green in PBS for 10 minutes at room temperature. Cells were washed 2 times in PBS to remove excess dye, and then resuspended in RPMI 1640 media. AlexaFluor647-labeled anti-hHER2 antibody (mouse IgG2a isotype targeting clinical Trastuzumab hHER2 epitope, Aduro Biotech) was serially diluted with MC38-hHER2 cells to achieve appropriate final concentrations for opsonization. MC38-hHER2 cells were coated in anti-hHER2 antibody dilutions 20 minutes prior to addition to BMDMs wells. When applicable, 18a antibody (SIRPα-blocking antibody, Aduro Biotech) was added directly to seeded BMDMs 20 minutes prior to MC38-hHER2 cell addition. 40,000 MC38-hHER2 cells (in 100 μL) were added to each well of 40,000 BMDMs to achieve a 1:1 ratio.

MC38-hHER2 cells were incubated with BMDMs for exactly 30 minutes. Wells were washed 2 times with PBS to remove excess MC38-hHER2 cells. Wells were imaged using the same microscope and automated technique as the target particle phagocytosis assay. Images were analyzed using a CellProfiler program that identified single BMDM cells and quantified the internalized MC38-hHER2 fluorescence intensity (CellTracker Green) within each BMDM cell. Average internalized fluorescence per cell was calculated per condition. Independent replicates were conducted on different days, and replicates were normalized to average eating for all conditions to control for day-to-day variation.

### Generation of cell lines

#### Design of synthetic receptors

The alpha helical SYNZIP Syn18-Syn17 binding pair was chosen due to its antiparallel orientation (Thompson et al., 2012). For each synthetic receptor design, the 41 amino acid sequence of Syn18 was placed N-terminally to one FNIII domain (Jacobs et al., 2012) to control the height above the membrane of the synthetic receptor protein (Figure S2a). The extracellular part was followed by the transmembrane domain of mouse SIRPα (Accession #P97797, amino acid 374-394), followed by intracellular regions of FcγRIIA (Accession #P12318, amino acid 240-317), CD3ζ (Accession #24161, amino acid 52-164), FcεRI γ-chain (Accession #P20491, amino acid 45-86), SIRPα (Accession #P97797, amino acid 395-513), CD22 (Accession #P35329, amino acid 722-862), PD-1 (Accession #Q02242, amino acid 191-288), and FcγRIIB (Accession #P08101, amino acid 232-330). A C-terminal GFP facilitated cell sorting and validated receptor localization. The entirety of each of the annotated intracellular domains was ordered as a separate gene fragment (Integrated DNA Technologies) and each complete receptor was cloned into pHR lentiviral expression vector (Clontech). Each region of the protein was amplified using PCR and fragments were combined using Gibson assembly.

#### Design of Syk and SHP1 sensors

mCh-Syk live phosphosensor was designed and utilized as previously published (Bakalar et al., 2018).

SHP1 interacts with phosphorylated SIRPα through interaction with its tandem SH2 domains. We created a sensor that specifically localizes to phosphorylated ITIMs by fusing the two SH2 domains of SHP1 to GFP to enable visualization of intracellular localization of the expressed protein. SH2 domain sequences were obtained from Addgene (pGEX-SHP-1(NC)-SH2, A #46496 (Machida et al., 2007)) and a linker region (GGGSGGGS) was placed between the SH2 domains and GFP. The sensor was cloned into pHR lentiviral expression vector under control of low-expression UBC promoter.

#### Generation of stable cell lines

HEK293T cells were grown to 70% confluency in a 6-well tissue-culture plate, and 160 ng pDM2.G (Clontech), 1.3 μg CMV 8.91 (Clontech), and 1.5 μg of target vector were transfected using TransIT-293T transfection reagent (Mirus Bio). Viral supernatant was collected 60 hours after transfection and spun at 1000 x g to remove HEK293T cells and used immediately. 1 mL of lentiviral supernatant was added to 5×10^5^ RAW 264.7 macrophages with 4 μg/mL of hexadimethrine bromide (Millipore) and cells were spinoculated at 300 x g for 30 minutes at room temperature. Cells were resuspended and plated into a 10 cm non-tissue-culture dish. Cells were sorted via fluorescence-activated cell sorting using an Aria Fusion cell sorter (Becton Dickinson) and the sorted population was expanded for use. For cell lines expressing synthetic receptors, function of Syn18 receptors was confirmed using AlexaFluor647-labeled soluble Syn17-F3L (see “Purification of Syn17-F3L”).

### Purification of Syn17-F3L

#### Design of Syn17-F3L

To create the Syn18 receptor ligand, the 42-amino acid sequence of Syn17 (Thompson et al., 2012) was followed by three repeats of the synthetic FNIII domain (Jacobs et al., 2012). Three Fibcon repeats were used to achieve a desired receptor-ligand interaction height. Finally, a C-terminal His-10 allowed for attachment to Ni-NTA lipids. Each region of the protein was amplified via PCR and fragments were combined using Gibson assembly into pET28 vector (Millipore) for expression in *E. coli*.

#### Syn17-F3L protein expression and purification

Syn17-F3L was expressed in Rosetta DE3 competent cells (Millipore). Cells were grown at 37°C to OD = 0.8 and expression was induced with 0.3 mM IPTG (Isopropyl b-D-1-thiogalactopyranoside, Calbiochem) overnight at 18°C. Cells were pelleted and resuspended in 25 mM HEPES pH 7.4 (4-(2-hydroxyethyl)-1-piperazineethanesulfonic acid, Thermo Fisher Scientific), 150 mM NaCl, 0.5 mM TCEP (tris(2-carboxyethyl)phosphine, Thermo Fisher Scientific) and 10 mM imidazole (Thermo Fisher Scientific), and lysed. After centrifugation, lysate was affinity purified using Cobalt-charged His-Trap Chelating column (GE Healthcare) through imidazole gradient elution. Peak fractions were gel-filtered using Superdex 200 column (GE Healthcare) into 25 mM HEPES pH 7.4, 150 mM NaCl, 0.5 mM TCEP. Protein was then labeled with AlexFluor647 NHS Ester (Thermo Fisher Scientific) as described in “Fluorescent labeling of proteins”.

### SLB TIRF imaging and quantification

#### Formation of planar SLBs

To image footprints of macrophages and GPMVs, planar SLBs were formed onto coverslips via fusion of SUVs to RCA-cleaned glass coverslips. PDMS (Polydimethylsiloxane, Sylgard) placed atop RCA-cleaned glass formed the imaging chamber, and 50 μL of MOPS buffer and 50 μL of non-fluorescent SUV solution were added. The SUVs were incubated for 20 minutes at room temperature. Excess SUVs were then removed by gently washing 5x with 60 μL of PBS. Appropriate antibody or CD47 dilutions were prepared, 60 μL were added to the washed SLB and incubated for 20 minutes. Excess protein was removed by gently washing the SLB 2x with PBS. The fluidity of proteins on the SLB was confirmed by using a spinning-disk confocal (Nikon) to examine diffusion of labeled molecules after photobleaching a small region of interest.

#### GPMV formation

GPMVs were made according to the protocol outlined by Sezgin et al (Sezgin et al., 2012). RAW 264.7 macrophages were seeded in a 6-well plate (Falcon). After adhering, they were rinsed with 1 mL GPMV buffer (10 nM HEPES (4-(2-hydroxyethyl)-1-piperazineethanesulfonic acid), 150 mM NaCl, 2 mM CaCl_2_, pH 7.4) before addition of 1 mL of vesiculation buffer (GPMV buffer plus 25 mM PFA (paraformaldehyde, Electron Microscopy Services) and 2 mM DTT (1,4-dithiothreitol, Anaspec). Cells were incubated for 1 hour at 37°C and GPMVs were collected by removing supernatant from cells.

#### TIRF imaging of GPMV and cell interfaces

Macrophages or GPMVs were added directly to planar SLBs containing AlexaFluor647-labeled anti-biotin IgG, AlexaFluor488-labeled CD47, or both. Cells or GPMVs were allowed to settle for 20 minutes prior to imaging. Engaged GPMVs or cells were located and identified using reflection interference contrast microscopy (RICM). Images were acquired using RICM and TIRF microscopy for all relevant fluorescent channels. When applicable, CD45 was labeled with 10 nM anti-CD45 antibody (clone 30-F11 Brilliant Violet 421, BioLegend).

#### Quantification of TIRF images

Images were quantified by identifying ROIs in RICM so as to not bias analysis by segmenting on a fluorescent channel. RICM images were flattened using a background (no cells or GPMV) image, then outlines of SLB-bound cells or GPMVs were identified. Fluorescence inside and outside these ROI outlines was measured and averaged for each fluorescent channel. The outside background measurement was subtracted from the average signal inside of the footprint. This was implemented using an automated custom macro in FIJI (Schindelin, J. et al., 2012). Data was processed and analyzed using Python 3.5 (python.org).

### Staining and quantification of phosphotyrosine

#### Fixation of macrophages with target particles

To stain for phosphotyrosine at macrophage-target interfaces, macrophages were seeded into 8-well imaging chambers with a coverslip glass bottom (Cellvis) and allowed to adhere for at 3-4 hours. If applicable, cells were treated with 3 μM SHP1/2 phosphatase inhibitor (NSC-87877, Sigma-Aldrich) for 20 minutes prior to target particle addition. Target particles coated in non-fluorescent SLBs plus AlexaFluor647-labeled anti-biotin IgG and/or AlexaFluor488-labeled CD47 were added to the seeded cells. After exactly 10 minutes at 37°C, cells were fixed for 10 minutes with 4% PFA in PBS. Cells were then permeabilized using 0.2% Tween-20 (Thermo Fisher Scientific) for 10 minutes and rinsed twice with PBS. Non-specific binding was blocked with 3% (w/v) bovine serum albumin (BSA, ChemCruz) in PBS with 0.5 g/mL Fc Block (BD Biosciences) for 30 minutes. Phosphotyrosine antibody (P-Y-1000 MultiMab, Cell Signaling Technology 8954S) was added to cells at 1:500 dilution in 1% BSA and incubated at room temperature for 1 hour. Cells were rinsed twice with 1% BSA, then incubated with a secondary antibody (AlexaFluor546-labeled goat anti-rabbit IgG, Invitrogen A11010) at a dilution of 1:1000 in 1% BSA for 1 hour at room temperature. Cells were washed twice with PBS before fluorescence and brightfield imaging with spinning-disk confocal microscope.

Interfaces between cells and bound (but not internalized) target particles were manually identified using brightfield images. Average pixel intensity within the interface was quantified using an automated custom macro in ImageJ.

#### Fixation and quantification of macrophages on SLB

To stain phosphotyrosine within footprints, macrophages were dropped onto a planar SLB and incubated at 37°C for 10 minutes. After exactly 10 minutes, cells were fixed and stained according to “Fixation of macrophages with beads.” Cells were imaged using RICM and TIRF imaging, and fluorescence within the footprint was measured as described in “Quantification of TIRF images.”

#### Fixation of BMDMs with tumor cells

To stain phosphotyrosine at BMDM-MC38 interfaces, 60,000 BMDMs were seeded into 8-well imaging chambers with a coverslip glass bottom. 40,000 MC38-hHER2 cells were incubated with various concentrations of AlexaFluor647-labeled anti-hHER2 antibody for 20 minutes at room temperature with continuous rotation. Opsonized MC38-hHER2 cells were added to BMDMs and incubated at 37°C for exactly 10 minutes. Cells were fixed and stained according to “Fixation of macrophages with beads.”

### Modeling

Activation and inhibition pathways were modeled using a set of 13 differential equations. Each equation represented a molecular species in the model, and the mass balance equations calculated species concentration change per unit time. Equations were implemented using Python 3.5 using *SciPy* software package functions for numerically solving ODEs (scipy.org). Simulations were run for 200 time points, a sufficient time to consistently reach steady state values for all species. Antibody concentrations were initiated from 1000 to 5000, and CD47 concentrations between 0 and 10000.

### Measurement of cell surface densities

BMDM, RAW 264.7, or MC38-hHER2 cells were resuspended at a density of 300,000 cells/mL in PBS with 5% FBS. On RAW 264.7 cells, Mouse Fc Block (rat anti-mouse CD16/CD32, BD Biosciences 553142) and p84 (provided by Aduro Biotech) were used to label FcγRs and SIRPα, respectively. On BMDMs, Fc Block and 18a with mutant Fc tail to prevent FcγR binding (provided by Aduro Biotech) were used to label FcγRs and SIRPα, respectively. On MC38-hHER2 cells, anti-hHER2 (mouse Fc Trastuzumab) and MIAP 301 (both provided by Aduro Biotech) were used to label hHER2 and CD47, respectively. Antibodies were labeled using AlexFluor647 NHS Ester (Thermo Fisher Scientific) (see “Fluorescent labeling of proteins”). Each antibody was added at a top concentration of 50 μg/mL, then serially diluted 4-fold into 8 sequential samples. This yielded a final range of 0.01 to 50 μg/mL antibody. Each antibody concentration was mixed with 30,000 cells and incubated at 4°C for 30 minutes to prevent internalization. Cells were washed twice by spinning for 5 minutes at 300 x g, removing supernatant, and replacing with PBS. Fluorescence of antibodies bound to cell surface was immediately measured on an Attune NxT flow cytometer. Surface antibodies were quantified using calibrated fluorescent beads.

### Statistical Analysis

All statistical analysis was performed in Python 3.5 using *SciPy* software package functions. The number of cells, particles, and interfaces quantified per experiment are indicated for each figure. In general, significance was based on a two-sample Student’s t-test for the mean values of experimental replicates, where *p<0.05, **p<0.01, and ***p<0.001.

## ACKNOWLEDGEMENTS

The authors would like to thank Fletcher Lab members, especially Carmen Chan and Aaron Joffe, for useful feedback and technical consultation; Cancer Research Laboratory Flow Cytometry Facility for cell sorting; Portnoy Lab of UC Berkeley for providing BMDMs. We are grateful to Aduro Biotech—specifically Andrea van Elsas, Meredith Leong, Sander van Duijnhoven, and Sanne Spijkers—for support, discussions, and supplying a range of reagents used in this study. This work was supported by the NIH R01 GM114671 (DAF), the Immunotherapeutics and Vaccine Research Initiative at UC Berkeley (DAF), the Miller Institute for Basic Research (DAF), and the Chan Zuckerberg Biohub (DAF). ECS was funded by an NSF-GRFP fellowship.

## AUTHOR CONTRIBUTIONS

ECS, EMS, and DAF contributed to study conception and design. ECS executed reconstitution experiments and modeling, and EMS executed tumor cell experiments and surface characterization. ECS and EMS analyzed and interpreted data. EV and BF provided reagents as well as scientific and therapeutic expertise. ECS wrote the manuscript with extensive input from EMS and DAF.

## DECLARATION OF INTERESTS

The authors declare no competing interests.

## Notes

### Competing Interest Statement

The authors have declared no competing interest.

