## Supplemental Figures and Tables for "Antibody:CD47 ratio regulates macrophage phagocytosis through competitive receptor phosphorylation"

### Supplemental Figure 1.

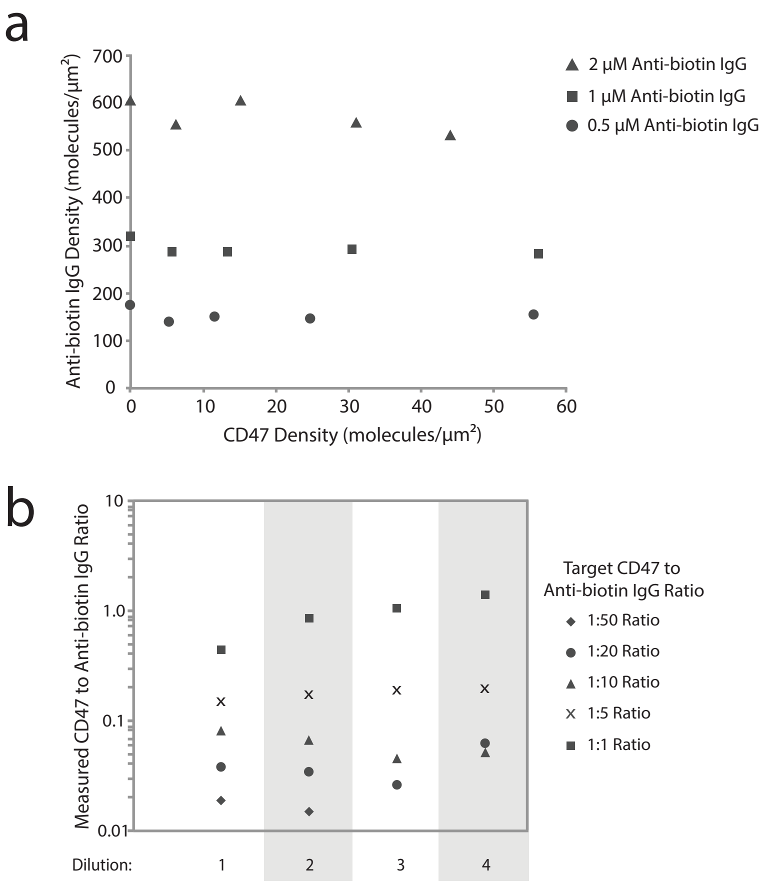

**Figure S1. Quantification of bead surface densities of antibody and CD47**

(a) Example target particle surface density data from phagocytosis assay summarized in Figure 1b. SLB-coated target particles were coated in different concentrations of AlexaFluor647-labeled anti-biotin IgG and AlexaFluor488-labeled CD47. Target particle fluorescence was measured via flow cytometry and compared to calibrated beads to calculate surface densities for each protein.

(b) Example target particle surface density data from phagocytosis assay summarized in Figure 1c. Target ratios of CD47 and anti-biotin IgG were created, then serially diluted 5-fold 3 times (for a total of 4 dilutions). Note that conditions that have no CD47 added or data points in which CD47 was diluted past detectable limits are not plotted.

### Supplemental Figure 2.

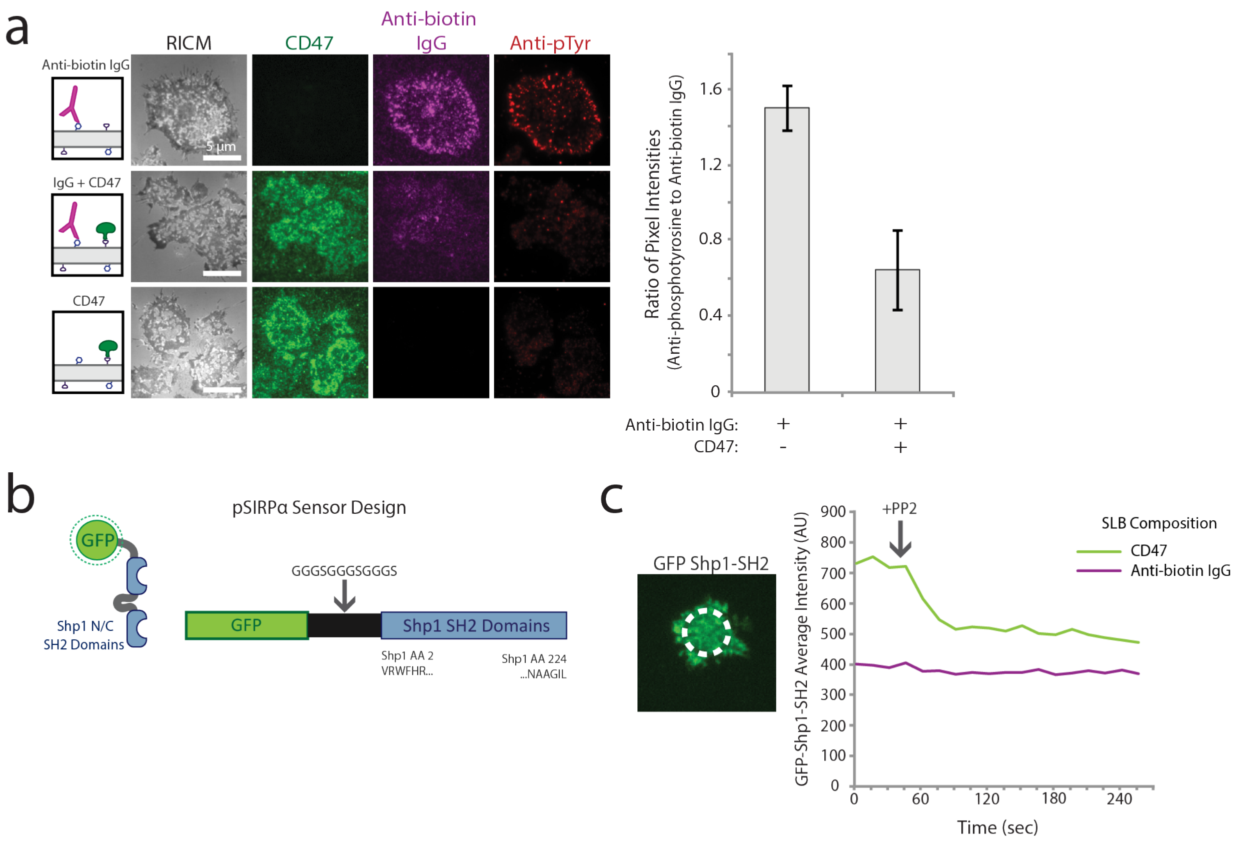

**Figure S2. Measurement and sensor of phosphorylation at immune interface**

(a) RAW 264.7 macrophages were dropped onto SLBs coated in AlexaFluor647-labeled anti-biotin IgG (magenta) with and without AlexaFluor488-labeled CD47 (green), and imaged using RICM and TIRF microscopy (left panel). The fluorescence ratio of anti-phosphotyrosine (red) to anti-biotin IgG was quantified for each footprint (right panel). Each condition is the average of >50 footprints. Bars represent mean ± s.e.m.

(b) Design of SHP1-GFP sensor for phosphorylated SIRPα. GFP was linked to the tandem SH2 domains of SHP1, which bind to phosphorylated ITIMs in the cytoplasmic tail of SIRPα.

(c) Effect of addition of 1μM Src family kinase inhibitor PP2 on SHP1-GFP sensor intensity. Src family kinase Lyn is responsible for phosphorylation of SIRPα, and decreasing SIRPα phosphorylation prevents SHP1 binding. Sensor intensity over time was measured in region indicated by white circle.

### Supplemental Figure 3.

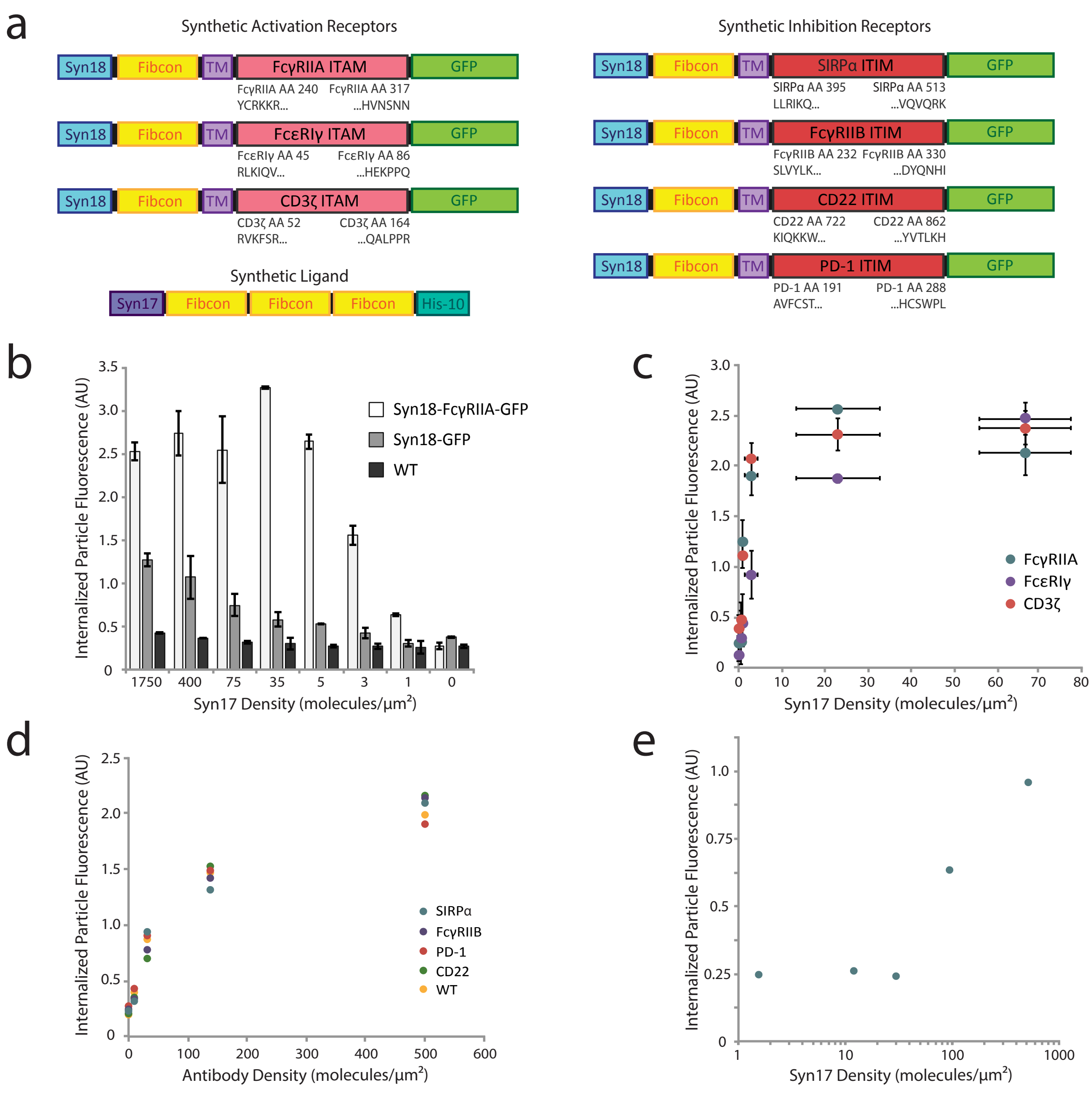

**Figure S3. Validation of ITAM- or ITIM-containing synthetic receptors**

(a) Schematic of synthetic chimeric receptor constructs. Synthetic activation receptors signaled via FcγRIIA, FcεRI γ-chain, and CD3 ζ-chain ITAM-containing intracellular domains. Synthetic inhibition receptors signaled via FcγRIIB, PD-1, SIRPα, and CD22 ITIM-containing intracellular domains.

(b) Target particles with varying surface densities of Syn17-F3L (0-1750 molecules/$\mu m^2$) were incubated with RAW 264.7 macrophages stably expressing Syn18-FcγRIIA-GFP receptor (light gray bars) and non-signaling Syn18-GFP (medium gray bars), as well as wild type macrophages (WT, dark gray bars). Average phagocytosis was quantified. Each condition is the average of three independent experiments representing a total of >300 cells. Bars represent mean ± s.e.m.

(c) Phagocytosis of Syn17-3L-coated target particles with macrophages expressing different Syn18 activating receptors. RAW 264.7 macrophages independently express Syn18 receptors containing FcγRIIA, FcεRI γ-chain, and CD3 ζ-chain intracellular signaling motifs. Target particles coated with increasing Syn17-F3L densities (0-75 molecules/$\mu m^2$) were simultaneously added to each receptor cell line, and average phagocytosis was quantified in the absence of an inhibitory ligand. Each condition is the average of three independent experiments representing a total of >300 cells. Dots represent mean ± s.e.m.

(d) Phagocytosis of anti-biotin IgG coated target particles by macrophages expressing different Syn18 inhibitory receptors. RAW 264.7 macrophages independently express Syn18 receptors containing FcγRIIB, PD-1, SIRPα, and CD22 intracellular signaling motifs. Target particles coated with increasing anti-biotin IgG surface densities (0-500 molecules/ $\mu m^2$) were simultaneously added to each receptor cell line as well as WT macrophages, and average phagocytosis was quantified in the absence of an inhibitory ligand. Each condition is the average of >100 cells.

(e) Phagocytosis of Syn17-F3L coated target particles by macrophages expressing Syn18-SIRPα-GFP.  Phagocytosis was quantified for target particles coated in increasing densities of Syn17-F3L (0-500 molecules/$\mu m^2$) in the absence of an activating ligand. Each condition is the average of >100 cells.

### Supplemental Figure 4.

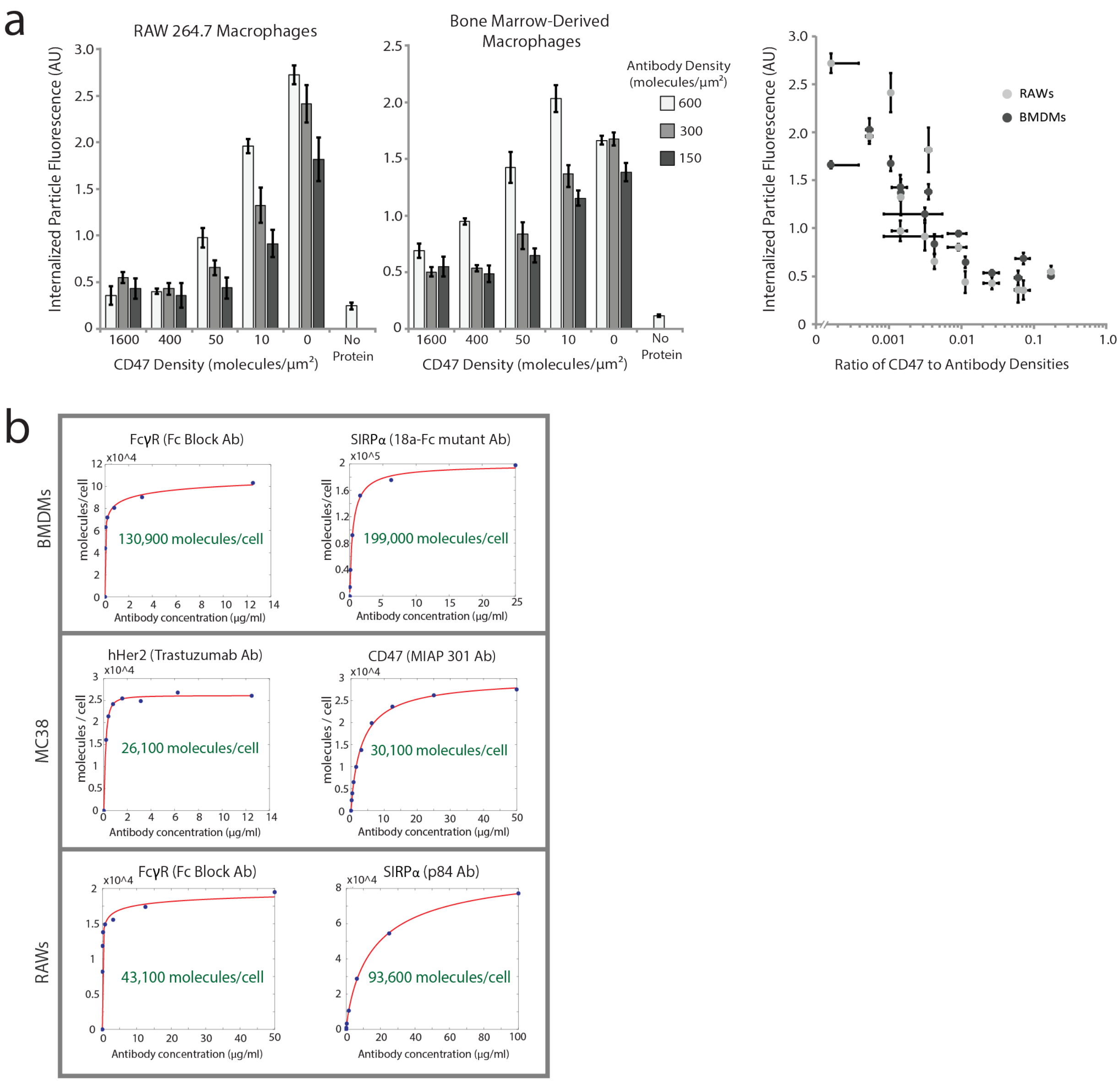

**Figure S4. Characterization of BMDM phagocytosis and cell surface densities**

(a) Phagocytosis comparison between RAW 264.7 macrophages and BMDMs. To create a panel of target particle compositions, three different anti-biotin IgG surface densities (150, 300, 600 molecules/$\mu m^2$) were combined with five different CD47 densities (0-1600 molecules/$\mu m^2$), plus an additional no protein (lipid only) control. All 16 different target particle densities were simultaneously added to RAWs and BMDMs, and average phagocytosis was quantified (left and center panels). Data replotted as a function of CD47:antibody ratio (right panel). Each condition is an average of 3 independent experiments representing >200 total cells. Bars represent mean ± s.e.m.

(b) Measurements of surface densities on RAW 264.7 macrophages, BMDMs, and MC38-hHER2 tumor cells. AlexaFluor647-labeled antibodies for FcγRs (Fc Block), SIRPα (18a Fc mutant and p84), hHER2 (anti-hHER2), and CD47 (MIAP 301) were increased to receptor saturating concentrations and measured via flow cytometry. Number of molecules per cell was estimated by fitting a Hill equation to each antibody titration and calculating $y_{max}$.

### Supplemental Table 1.

| Equations implemented in interacting kinetic model ($p$ denotes phosphorylated state) |
|---|
| $\frac{\partial}{\partial t}[FcR] = -k_1[Ab][FcR] + k_2[FcR-Ab]$ |
| $\frac{\partial}{\partial t}[FcR-Ab] = k_1[Ab][FcR] - k_2[FcR-Ab] - k_3[FcR-Ab] + k_{13}[SIRP\alpha - Shp1 - FcR_p]$ |
| $\frac{\partial}{\partial t}[Ab] = -k_1[Ab][FcR] + k_2[FcR-Ab]$ |
| $\frac{\partial}{\partial t}[FcR_p] = k_3[FcR-Ab] - k_4[FcR_p][Syk] + k_5[FcR-Syk] - k_{11}[SIRP\alpha - Shp1][FcR_p] + k_{12}[SIRP\alpha - Shp1 - FcR_p]$ |
| $\frac{\partial}{\partial t}[FcR-Syk] = k_4[FcR_p][Syk] - k_5[FcR-Syk]$ |
| $\frac{\partial}{\partial t}[Syk] = -k_4[FcR_p][Syk] + k_5[FcR-Syk]$ |
| $\frac{\partial}{\partial t}[SIRP\alpha] = -k_6[SIRP\alpha][CD47] + k_7[SIRP\alpha - CD47]$ |
| $\frac{\partial}{\partial t}[CD47] = -k_6[SIRP\alpha][CD47] + k_7[SIRP\alpha - CD47]$ |
| $\frac{\partial}{\partial t}[SIRP\alpha - CD47] = k_6[SIRP\alpha][CD47] - k_7[SIRP\alpha - CD47] - k_8[SIRP\alpha - CD47]$ |
| $\frac{\partial}{\partial t}[SIRP\alpha_p] = k_8[SIRP\alpha - CD47] - k_9[SIRP\alpha_p][Shp1] + k_{10}[SIRP\alpha - Shp1]$ |
| $\frac{\partial}{\partial t}[SIRP\alpha - Shp1] = k_9[SIRP\alpha_p][Shp1] - k_{10}[SIRP\alpha - Shp1] - k_{11}[SIRP\alpha - Shp1][FcR_p] + k_{12}[SIRP\alpha - Shp1 - FcR_p] + k_{13}[SIRP\alpha - Shp1 - FcR_p]$ |
| $\frac{\partial}{\partial t}[Shp1] = -k_9[Shp1][SIRP\alpha_p] + k_{10}[SIRP\alpha - Shp1]$ |
| $\frac{\partial}{\partial t}[SIRP\alpha - Shp1 - FcR_p] = k_{11}[SIRP\alpha - Shp1][FcR_p] - k_{12}[SIRP\alpha - Shp1 - FcR_p] - k_{13}[SIRP\alpha - Shp1 - FcR_p]$ |

**Table S1. Differential equations implemented for interacting kinetic model**

Equations were simulated in Python 3.5.

### Supplemental Table 2.

| Rate Constant | Value | Citation | Notes |
|---|---|---|---|
| $k_1$ | $8.2 \times 10^3$ $M^{-1}s^{-1}$ | Li et al, 2007 | |
| $k_2$ | $5.7 \times 10^{-3}$ $s^{-1}$ | Li et al, 2007 | |
| $k_3$ | 0.15 $s^{-1}$ | Barua et al, 2012 | Estimated from value for unphosphorylated Lyn |
| $k_4$ | $1.8 \times 10^5$ $M^{-1}s^{-1}$ | Barua et al, 2012 | |
| $k_5$ | 0.3 $s^{-1}$ | Barua et al, 2012 | |
| $k_6$ | $2 \times 10^5$ $M^{-1}s^{-1}$ | Brooke et al, 2004 | |
| $k_7$ | 5.3 $s^{-1}$ | Brooke et al, 2004 | |
| $k_8$ | 0.15 $s^{-1}$ | Barua et al, 2012 | Same as $k_3$; Assumes no ITAM vs. ITIM preference |
| $k_9$ | $1.8 \times 10^6$ $M^{-1}s^{-1}$ | Barua et al, 2012 | Estimate from other SFKs |
| $k_{10}$ | 0.1 $s^{-1}$ | Barua et al, 2012 | |
| $k_{11}$ | $1.8 \times 10^5$ $M^{-1}s^{-1}$ | Ren et al, 2011 | Estimated from $k_d$ value |
| $k_{12}$ | 5 $s^{-1}$ | Ren et al, 2011 | |
| $k_{13}$ | 66 $s^{-1}$ | Selner et al, 2014 | Measured value for peptide w/I at Y -2 position |

**Table S2. Rate constants utilized in phagocytosis kinetic model**

Constants that were estimated from published values are noted.
